# Epitranscriptomic regulation of the response to the air pollutant naphthalene in mouse lungs: from the perspectives of specialized translation and tolerance linked to the writer ALKBH8

**DOI:** 10.1101/727909

**Authors:** Andrea Leonardi, Nataliia Kovalchuk, Lei Yin, Lauren Endres, Sara Evke, Steven Nevins, Samuel Martin, Peter C. Dedon, J. Andres Melendez, Laura Van Winkle, Qing-Yu Zhang, Xinxin Ding, Thomas J. Begley

## Abstract

**Background:** The epitranscriptomic writer Alkylation Repair Homolog 8 (ALKBH8) is a tRNA methyltransferase that modifies the wobble uridine of selenocysteine tRNA to promote the specialized translation, via stop codon recoding, of proteins that contain selenocysteine. Corresponding selenoproteins play critical roles in protecting against reactive oxygen species and environmental stress. Using a novel animal model deficient in *Alkbh8*, we have investigated the importance of epitranscriptomic systems in the response to naphthalene (NA), an abundant polycyclic aromatic hydrocarbon, glutathione depleter and lung toxicant found in tobacco smoke, gasoline and mothballs.

**Objectives:** Our goal was to define the molecular reprogramming of Alkbh8 deficient (*Alkbh8^def^)* mice and evaluate the roles that the epitranscriptomic writer ALKBH8 and selenoproteins play in mitigating NA-induced toxicity and lung dysfunction.

**Methods:** We performed basal lung analysis and NA exposure studies using WT, *Alkbh8^de^*^f^ and *Cyp2abfgs-null* mice, the latter of which lack the cytochrome P450 enzymes required for NA bioactivation. We characterized gene expression, molecular markers of damage, viability and tolerance to NA.

**Results:** Under basal conditions, lungs from *Alkbh8^def^* mice have increased oxidation-reduction potential (ORP) and 8-isoprostane levels, and have reprogrammed at the molecular level to display increased stress response transcripts. In addition, the ALKBH8 writer deficient lungs have a decreased GSH/GSSG ratio. *Alkbh8^def^* mice are more sensitive to NA than WT, showing higher susceptibility to lung damage both at the cellular and molecular levels. WT mice develop a tolerance to NA after 3 days, defined as resistance to a high challenging dose after repeated exposures, which is absent in *Alkbh8^def^* mice, with writer deficient not surviving NA exposure.

**Discussion:** We conclude that the epitranscriptomic writer ALKBH8 plays a protective role against NA-induced lung dysfunction and promotes NA tolerance. Our work provides an early example of how epitranscriptomic systems can regulate the response to environmental stress *in vivo*.

## 1. Introduction

Epitranscriptomic marks in the form of RNA modifications are catalyzed by specific enzymes or RNA-protein complexes on the base or ribose sugar of the canonical adenosine, cytosine, guanosine and uridine nucleosides. RNA modifications can include simple changes such as methylation or acetylation, or hyper modifications involving multiple enzymes and cofactors. Transfer RNA (tRNA) is the most heavily modified RNA species with an average of 13 RNA modifications comprising ∼17% of the 80-90 ribonucleosides. Epitranscriptomic marks can be located throughout tRNA molecules and can affect tRNA stability, folding, localization, transport, processing, and function (Jackman & Alfonzo, 2013). Modifications in the anticodon loop of tRNA can function as regulators of translation by promoting or preventing codon-anticodon interaction, by altering charge or base-pairing potential. Global studies using cell systems have demonstrated that anticodon specific epitranscriptomic marks, as well as others, are dynamically regulated during cellular stress. Further it has been shown that the collection of epitranscriptomic marks change depending on the mechanism of action of the toxicants (C. T. Y. Chan et al., 2015, 2010; Dewe, Fuller, Lentini, Kellner, & Fu, 2017).

Toxicant specific reprogramming of the epitranscriptome housed on tRNA has been demonstrated in bacterial, yeast and human cell studies (C. T. Y. Chan et al., 2015, 2010; Dewe et al., 2017). For example, in response to oxidizing agents such as γ-radiation, hydrogen peroxide (H_2_O_2_), tert-Butyl hydroperoxide (TBHP) or peroxynitrile (ONOO^-^), there is a significant increase in m^5^C, ncm^5^U, and i^6^A levels in yeast tRNA while the same modifications are unaffected in response to alkylating agents. In contrast, Um, mcm^5^U, mcm^5^s^2^U, m^2^G, and m^1^A modification levels increase in response to alkylating agents ethylmethanesulfonate (EMS), methylmethanesulfonate (MMS), isopropyl methanesulfonate (IMS) and N-methyl-N’-nitro-N-mitrosoguanidine (MNNG), and are conversely unaffected by treatment with oxidizing agents. These data support the idea that there is a highly predictive and toxicant-dependent system of translational control used to respond to cellular stress. Specifically the reprogrammed tRNA promotes the selective translation of codon-biased mRNAs in order to translate critical survival proteins. Codon biased transcripts being more efficiently translated by increased epitranscriptomic marks has been observed in bacteria (Chionh et al., 2016), yeast (Bauer & Hermand, 2012; Bauer et al., 2012; C. T. Y. Chan et al., 2012; Deng et al., 2015), mouse embryonic fibroblasts (Endres et al., 2015) and human melanoma cells (Rapino et al., 2018). A specific example of epitranscriptomic marks driving codon biased translation is found with the RNA modifications 5-methoxycarbonylmethyluridine (mcm^5^U) and 5-methoxycarbonylmethyluridine-2’-O-methyluridine (mcm^5^Um) housed on wobble uridine of selenocysteine tRNA (tRNA^Sec^) in mice and humans. The epitranscriptomic marks mcm^5^U and mcm^5^Um are an integral part of the specialized addition of the selenocysteine (Sec) amino acid into proteins. The resulting selenoproteins are made by stop codon recoding (Copeland, 2003; Copeland, Fletcher, Carlson, Hatfield, & Driscoll, 2000; Endres et al., 2015; Low & Berry, 1996; Songe-Moller et al., 2010), where mcm^5^U and mcm^5^Um promote binding of the tRNA^Sec^ anticodon to the UGA stop codon on mRNA. Corresponding selenoprotein transcripts are inherently codon-biased because of this unusual in-frame UGA stop codon as well as the actual stop codon(Doyle et al., 2016). Importantly though, other core elements are required for stop codon recoding and they include a stem-loop structure in the 3’ untranslated region (3’UTR) of the transcript termed the <u>Sec</u> Insertion <u>S</u>equence (SECIS) element, a SECIS Binding Protein (SBP2), and SEC-specific elongation factors (EFSec) (Driscoll & Copeland, 2003).

Mammalian Alkylation Repair Homolog 8 (ALKBH8) is a writer enzyme that is required for the formation of modified wobble uridines on tRNA^Sec^ to promote UGA stop codon recoding (Dragony Fu et al., 2010; Songe-Moller et al., 2010). ALKBH8 has three functional domains, an RNA recognition motif (RRM), a 2-oxogluterate Fe^2+^-dependent oxygenase (2-OG-Fe(II)) domain, sometimes referred to as the AlkB domain, and a methyltransferase domain (Fig. 1). The Elongator complex which includes six proteins (Elp1-6) is required for the carboxymethylation of uridine (U) to 5-carbonylmethyluridine (cm^5^U) (Bauer & Hermand, 2012; Bauer et al., 2012; Karlsborn et al., 2014). ALKBH8 methyltransferase activity on the wobble uridine of tRNA^Sec^ uses 5-carbonylmethyluridine (cm^5^U) as a substrate to produce mcm^5^U. ALKBH8 also catalyzes wobble mcm^5^U in tRNA^Lys(UUU)^, tRNA^Gln(UUG)^ and tRNA^Glu(UUC)^ ^and^ tRNA^Arg(UCU)^(D. Fu et al., 2010; Hatfield & Gladyshev, 2002; Songe-Moller et al., 2010; Sprinzl & Vassilenko, 2005). Furthermore, mcm^5^U can undergo additional modifications to mcm^5^Um (tRNA^Sec^)(Endres et al., 2015; Songe-Moller et al., 2010), mcm^5^s^2^U (tRNA^Glu(UUC,^ tRNA^Arg(UCU)^ ^and^ tRNA^Lys(UUU)^)(Sprinzl & Vassilenko, 2005), and mchm^5^U (tRNA^Gly(UCC)^ and tRNA^Arg(UCG)^)(Van Den Born et al., 2011). Among these further modifications to mcm^5^U, only mchm^5^U is known to be catalyzed directly by ALKBH8 via its 2OG-Fe(II) oxygenase domain(Van Den Born et al., 2011). The mcm^5^s^2^U modification depends on the CTU1/CTU2 enzyme complex, while formation of mcm^5^Um is dependent on ALKBH8 (Dewez et al., 2008; Endres et al., 2015; Songe-Moller et al., 2010).

**Figure 1.**
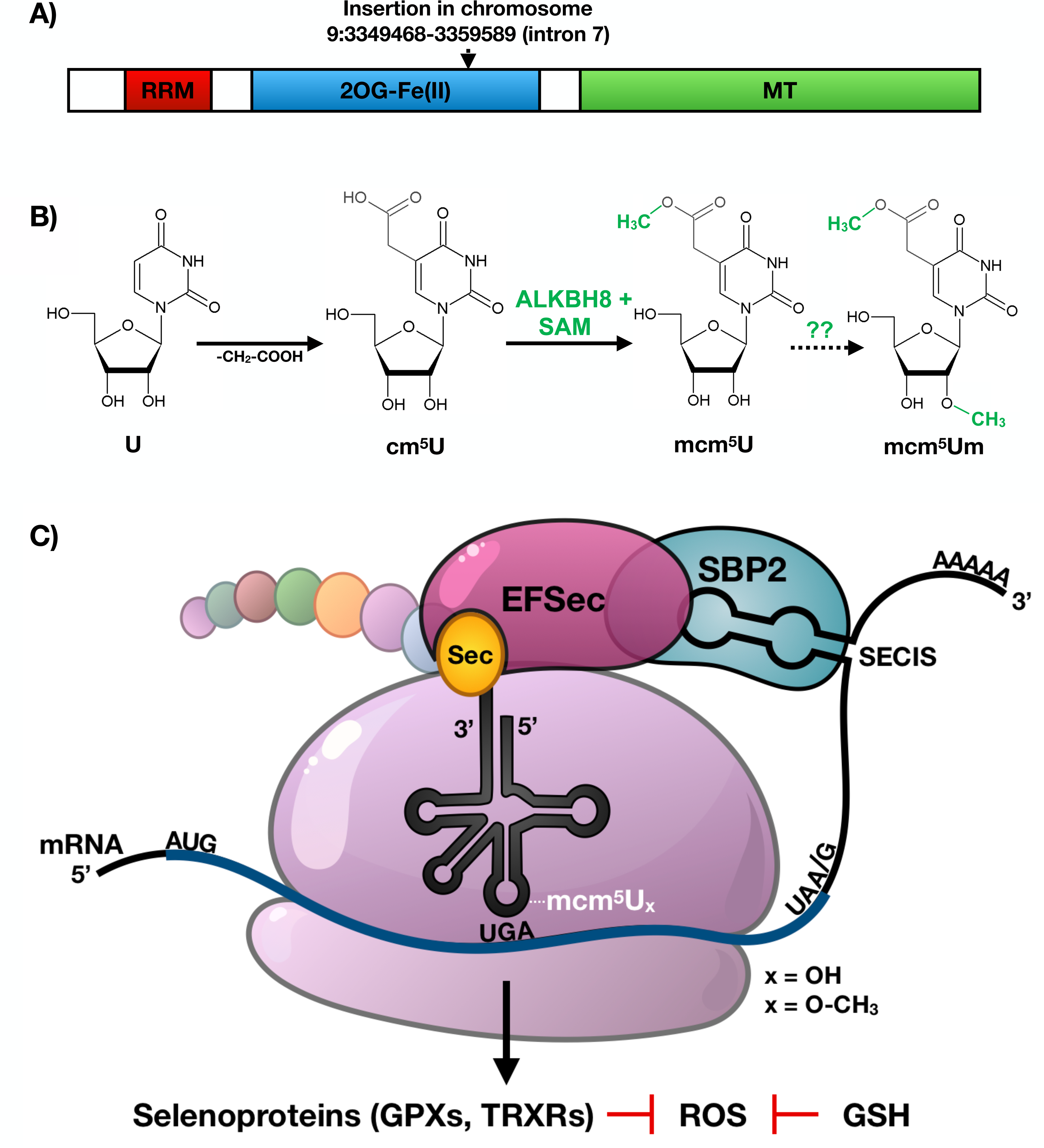
ALKBH8 domain structure, chemistry and role in translation of selenoproteins. **A)** The ALKBH8 protein contains RNA recognition motif (RRM), 2-oxogluterate and Fe^2+^-dependent oxygenase (2OG-Fe(II)), and methyltransferase (MT) domains. Included is the insertion area of the *β-galactosidase/neomycin* resistance cassette (Stryke et al., 2003). **B)** Formation of mcm^5^U and mcm^5^Um. Uridine (U) is first carboxymethylated to 5-carbonyl-methyluridine (cm^5^U) by elongator (ELP) proteins. Next, ALKBH8 catalyzes the formation mcm^5^U using S-adenosyl methionine (SAM) as a methyl donor. ALKBH8 is also required for the formation of mcm^5^Um. **C)** Selenoprotein synthesis requires a specialized mechanism of translation. In addition to a modified tRNA^Sec^, elongation factors, SECIS Binding Protein 2 (SPB2), and 3’ UTR Selenocysteine Insertion Sequence (SECIS) elements are required for translation of the UGA stop codon found in transcripts corresponding to selenocysteine-containing proteins.

The mcm^5^U and mcm^5^Um modifications on tRNA^Sec^ help translationally regulate the production of selenoproteins, which comprise an important group of activities involved in development, reproduction and protection against oxidative stress. Selenoproteins such as glutathione peroxidase (GPX) family members catalyze the reduction of hydrogen peroxide and lipid peroxides by glutathione and play an essential role in protecting cells against oxidative damage. Thioredoxin reductases (TRXRs) are involved in maintaining the reduced state of thioredoxins (TRXs), which function to reduce oxidative stress and play critical roles in development.

We have previously shown that deficiency in ALKBH8 prevents a H_2_O_2_ induced increase in mcm^5^Um and Alkbh8 deficient mouse embryonic fibroblasts (MEFs) have decreased selenoprotein expression levels, increased ROS and show sensitivity to the ROS-producing agent H_2_O_2_(Endres et al., 2015). ALKBH8 deficient mice are viable under normal care conditions which include a 12-hour light/dark cycle, 68-72°C temperature controlled environment, a standardized and balanced diet with free access to water, and minimal stress-inducing noise, odors and handling. Their ability to thrive in a normal environment suggests that *Alkbh8^def^* mice adapt to the epitranscriptomic deficiency. Similar to other important stress and selenoprotein deficiency (Haan et al., 1998), the ALKBH8 deficient mice may need to experience an external stress to display observable phenotypes.

Naphthalene (NA) is a polyaromatic hydrocarbon (PAH) and exposure to this environmental toxicant can promote ROS, DNA damage and disrupt cellular glutathione levels (ATSDR, 2005; D. Bagchi, Bagchi, Balmoori, Vuchetich, & Stohs, 1998; M. Bagchi, Bagchi, Balmoori, Ye, & Stohs, 1998; Carratt et al., 2019; Lin et al., 2005; Plopper et al., 2013a). NA is found in cigarette smoke, gasoline, and mothballs (Jia & Batterman, 2010; Kakareka & Kukharchyk, 2003; Sudakin, Stone, & Power, 2011) as well as being derived in a number of industrial processes. We postulated that ALKBH8 dependent epitranscriptomic marks would play an important role in response to the ubiquitous environmental toxicant NA. Notably there are few animal studies testing the importance of epitranscriptomic systems in response to PAHs. In mouse, rat and human respiratory tissue, cytochrome P450 (CYP) enzymes bioactivate NA to reactive and toxic metabolites. NA exposure is highly prevalent in the human population as detectable metabolite (1- and 2-hydroxynaphthalene) levels were found in urine of every one of 2,749 human civilians tested across 30 U.S. cities. In addition to its 100% detection rate in urine, NA contribute approximately 75% to the total concentration of all PAHs tested in the study (Z. Li et al., 2008). NA is currently classified as a possible human carcinogen(IARC, 2002) and has been shown to cause lesions and tumors in rodents, although the mechanism of toxicity and carcinogenicity is still under debate(Abdo et al., 1992; Abdo, Grumbein, Chou, & Herbert, 2001). Notably NA has been shown to cause oxidative DNA damage(Lin et al., 2005), and DNA adduct formation(Buchholz et al., 2019; Carratt et al., 2019). All tumor formation in both mice and rats was accompanied by cytotoxicity, with recurrent damage and repair/proliferation thought to be one of the driving forces behind NA-induced tumor formation in the lung (DeStefano-Shields, Morin, & Buckpitt, 2010; Van Winkle, Buckpitt, Nishio, Isaac, & Plopper, 1995). Importantly, glutathione (GSH) depletion is a major determinant in NA-induced lung toxicity, and precedes observable injury. Mice treated with the GSH depleter dimethylmaleate before exposure suffered significantly more injury to the nasal and intrapulmonary airway epithelium(Phimister, Lee, Morin, Buckpitt, & Plopper, 2004) and mice treated with the GSH synthesis inhibitor buthionine sulfoximine exhibited similar effects(West et al., 2003, 2002). Treatment of mice with glutathione prodrugs prevent lung cell toxicity and necrosis(Phimister, Nagasawa, Buckpitt, & Plopper, 2005). Selenoprotein H plays a role in glutathione synthesis by acting as a transcription factor to promote transcription of genes critical for glutathione synthesis as well as exhibiting oxidoreductase activity(Panee, Stoytcheva, Liu, & Berry, 2007). As such, disrupted ALKBH8 epitranscriptomic systems leading to dysregulated translation of selenoproteins required for GSH synthesis, use and redox balance are likely to sensitize mice to NA.

In the following study we have tested the hypothesis that the ALKBH8 epitranscriptomic writer plays a protective role against NA-induced lung toxicity *in vivo*. We have characterized the lungs of *Alkbh8* deficient (*Alkbh8^def^*) mice under normal conditions to elucidate how these mice adapt to an epitranscriptomic defect. We have also performed NA exposure studies on WT, A*lkbh8^def^*, and *Cyp2abfgs-null* mice. The *Cyp2abfgs-null* mice are a total-body knockout of a 12-gene cluster of lung preferential CYPs needed for NA bioactivation in the lung (Lei Li, Megaraj, Wei, & Ding, 2014). As NA bioactivation is a well-documented requirement of NA toxicity as previously described, the *Cyp2abfgs-null* mice served as our negative controls (L. Li et al., 2011; Lei Li et al., 2017).

Under basal conditions, lungs from the writer deficient mice have increased oxidation reduction potential (ORP), 8-isoprostane levels, and have reprogrammed at the molecular level to display increased stress response transcripts. In addition, the ALKBH8 writer deficient lungs have decreased GSH and increased GSSG, resulting in a disrupted GSH/GSSG ratio which we predict is a direct result of increased oxidative stress. *Alkbh8^def^* mice are significantly more sensitive to NA than WT, showing higher susceptibility to lung damage both at the cellular and molecular levels. Tolerance to NA is defined as resistance to a high challenging dose after repeated exposures. WT mice develop tolerance to NA after 3 days while *Alkbh8^def^* mice fail to develop tolerance to NA and die. Our data supports a model in which the ALKBH8 writer and epitranscriptomic marks allow cells to develop tolerance and adapt to chronic NA stress. Our work highlights that epitranscriptomic systems can regulate the response to environmental stress *in vivo*.

## Material and Methods

### 2.1. Animal Experiments

This study was carried out in strict accordance with the recommendations in the guide for the Care and Use of Laboratory Animals of the National Institutes of Health. The protocol was approved by the University at Albany Institutional Animal Care and Use Committee (Albany, NY) Protocol #17-016 for breeding and basal characterization and #18-011 for naphthalene challenge. Additionally, it was approved by the Wadsworth Center IACUC committee protocol 14-331 (Albany, NY) and The University of Arizona IACUC, protocol 17-335 (Tucson, AZ). Four homozygous strains of C57BL/6 mice were bred: Wild-type (WT), *Alkbh8* deficient (*Alkbh8^de^*^f^) and *Cyp2abfgs-null*. Wild-type mice were purchased from Taconic Biosciences (Rensselaer, NY). *Alkbh8^de^*^f^ mice were created using an insertional mutagenesis approach targeting the *Alkbh8* gene in the parental E14Tg2a.4 129P2 embryonic stem cell line using a vector containing a splice acceptor sequence upstream of a *β-geo* cassette (β-galactosidase/neomycin phosphotransferase fusion), which was inserted into intron 7 at chromosome position 9:3349468-3359589, creating a fusion transcript of *Alkbh8-β-geo* (Stryke et al., 2003), as previously described (Endres et al., 2015). Multiplex qRT-PCR and relative cycle threshold analysis (ΔΔ CT) on genomic DNA derived from tail biopsies was used to determine animal zygosity for *Alkbh8^def^* with TaqMan oligonucleosides specific for neomycin (*Neo*, target allele) and T-cell receptor delta (*Tcrd*, endogenous control). Neo forward primer (5’-CCA TTC GAC CAC CAA GCG-3’), Neo reverse primer (5’-AAG ACC GGC TTC CAT CCG-3’), Neo Probe (5’-FAM AAC ATC GCA TCG AGC GAG CAC GT TAMRA-3’), Tcrd forward primer (5’-CAG ACT GGT TAT CTG CAA AGC AA-3’), Tcrd reverse primer (5’-TCT ATG CAA GTT CCA AAA AAC ATC-3’), and Tcrd probe (5’-VIC ATT ATA ACG TGC TCC TGG GAC ACC C TAMRA -3’). *Cyp2abfgs-null* mice were generated using the *in vivo* Cre-mediated gene deletion approach as described in (Lei Li et al., 2014), in which a 1.4-megabase pair genomic fragment containing 12 *Cyp* genes is deleted, including *Cyp2a, Cyp2b, Cyp2f, Cyp2g*, and *Cyp2s* subfamilies. Genotyping of *Cyp2abfgs-null* mice was performed as described in (Lei Li et al., 2014). A double mutant strain was established via a cross of homozygous *Cyp2abfgs-null <u>x</u> Alkbh8^def^* mice creating *Cyp2abfgs-null/Alkbh8^def^* double mutants (denoted as *DM* in this manuscript).

Male mice in age between 8-12 weeks were used in all experiments. Mice were sacrificed via CO_2_ asphyxiation and lungs excised, then flash-frozen and stored at -80°C, or immediately processed for downstream experiments.

### 2.2. Naphthalene solution preparation and I.P. injection

NA (99% pure) and corn oil were purchased from Sigma-Aldrich (St. Louis, MO). Naphthalene was dissolved in corn oil for a concentration of 20 mg/ml for acute doses or 80 mg/ml for subchronic doses. The concentrations were optimized to ensure accurate NA dosing while minimizing corn oil-induced effects. Due to the volatile nature of NA, solutions were stored at -80°C and made fresh at the start of each experiment. In the case of multiple injections, solutions were stored at -80°C and discarded after 5 days. For injections, mice were weighed, placed under brief anesthesia (∼1 minute) using 5% isofluorane gas, manually restrained, injection site wiped with 70% ethanol, and the corn oil or NA solution injected into the intraperitoneal cavity in the lower left abdominal quadrant using Coviden™ Monoject™ 27g, ½ inch standard hypodermic needles (Fisher Scientific, Franklin, MA) and sterile 1ml Coviden™ Monoject™ tuberculin syringes (Fisher Scientific) with 50-200 mg/kg of NA or an equal volume of corn oil proportional to body weight. In case of multiple doses, the injections were administered between 10 and 11 AM (US EST) 24h apart. Mice were observed for recovery from anesthesia, which typically occurred within a few seconds of placing the mouse back in the cage. Mice were further monitored for signs of excess stress and weight loss. Mice were sacrificed 24 h after injection, or 24 h after last injection in case of multiple doses.

### 2.3. RNA extraction and Quantitative real-time PCR (qRT-PCR)

Total RNA was isolated from mouse lungs under basal and NA-challenged conditions. Immediately after excision, fresh lungs were placed in Trizol reagent (Invitrogen, CA), cut into small pieces with scissors, and homogenized in 2 ml Trizol with a Tissue-Tearor homogenizer (BioSpec, Bartlesville, OK). Samples were left at room temperature for 10 minutes then placed on dry ice and transferred to -80°C for temporary storage. RNA isolation was performed using the standard Trizol reagent manufacturer’s protocol (http://tools.thermofisher.com/content/sfs/manuals/trizol_reagent.pdf). *Alkbh8* transcript levels were measured in the lung using a Taqman Gene Expression assays. All qRT-PCR was carried out using a Thermo Fisher Scientific QuantStudio™ 3 Real-Time PCR system (Waltham, MA). Each sample was tested in triplicate and normalized to a *β-Actin* endogenous control. Statistically significant differences between the ΔΔCT values were determined using an unpaired Student’s t-test. Error bars denote standard error of the mean.

#### 2.3.1 Targeted mRNA sequencing

Basal, 24-hour corn oil, and 24-hour naphthalene treated WT and *Alkbh8^de^*^f^ lung RNA was extracted using the QIAGEN RNease mini kit (Qiagen, Germantown, MD). Samples were sent to Qiagen Genomic Services (Frederick, MD) for quality control (QC), library prep and targeted RNA sequencing. RNA integrity and concentration were evaluated using Agilent TapeStation 4200 and Thermo Scientific Nanodrop spectrophotometer, respectively. The starting RNA input for library prep was 500 ng. QIAseq Targeted RNA with a custom panel of 350 stress response transcripts specific to DNA damage, oxidative stress, protein damage, inflammation, metabolism and signaling pathways were targeted for library prep and sequencing. The targeted libraries were pooled together and sequenced using miSeq™ 500 system (Illumina, San Diego, CA). See Supplemental Table 1 for complete gene list. Samples were processed in duplicate (2 mouse lungs per condition, per genotype). FASTQ files for each sample were obtained from BaseSpace and were further analyzed using the QIAGEN GeneGlobe Data Analysis Center (QIAGEN.com). Linear fold-change values were reported after normalization to WT (basal conditions) and WT corn oil treated (NA challenged conditions) samples, respectively, and assembled into a heat map using Java TreeView (Supplemental Fig. 1).

### 2.3. Measurements of Oxidation Reduction Potential (ORP)

Lung tissue was solubilized in extraction buffer (0.1 g per 1 mL phosphate buffered saline + 0.1% v/v Triton-X-100) using a rotor/stator tissue homogenizer. 30 mL of lysate was then applied to a filter-based sensor inserted into the RedoxSYS® galvanostat (RedoxSYS, Aytu BioScience, Inc., Englewood, CO). An electrochemical circuit was made when the sample (drawn by liquid flow) entered the sensor cell, after which an initial ORP measurement was made followed by the application of a current sweep through the sample, which exhausts the antioxidants present in the sample; the amount of current required to do this was determined over a 3-minute period. The primary output was a measurement of static ORP (mV), also known as the redox potential, representing the potential for electrons to move from one chemical species to another.

### 2.4. Histopathology

WT, *Alkbh8^def^*, *Cyp2abfgs-null* and *Alkbh8^def^/Cyp2abfgs-null* double mutant mice were exposed to NA as described above and sacrificed 24h after last injection by CO_2_ asphyxation. Lungs and trachea were exposed, a small incision was placed in the trachea, and the lungs were slowly inflated with 0.8 mL Z-Fix formalin based fixative (Anatech LTD, Battle Creek, MI) using a sterile, disposable blunt end needle (22 gauge, 0.5 in, Part #B22-50, SAI Infusion Technologies, Lake Villa, IL). The lungs were placed into a 50 ml microcentrifuge tube containing an additional 20 ml Z-fix. After 24h, the fixative was removed and replaced with 70% ethanol. Samples were sent to either the Wadsworth histopathology core (Albany, NY) or Mass Histology Service, Inc (Worcester, MA) for processing, paraffin embedding, sectioning and H&E staining. Mass Histology Service, Inc. captured 40X slide scan images of lung airways and alveolar regions. H&E stained slides were analyzed by board-certified pathologist at Mass Histology Service, Lawrence McGill, DVM, PhD, DACVP, and assessed for pathologies and extent of damage, with specific focus on club cells, under basal, NA or corn oil conditions.

### 2.5. DNA Damage Response analysis via γ-H2AX foci staining on primary lung cells

3 WT and 3 *Alkbh8^def^* mice under basal conditions were sacrificed using CO_2_ asphyxiation. Lungs and trachea were exposed, a small incision was placed in the trachea, and the lungs slowly inflated with 250 U/ml collagenase I (Product # C9891, Sigma-Aldrich), 4.3 U/ml lyophilized elastase (Catalog # LS002292, Worthington-Biochem, Lakewood, NJ), and 0.006% deoxyribonuclease I (Catalog #: LS002139, Worthington-Biochem, Lakewood, NJ) in DMEM/F-12 media (Catalog# 11320033, Thermo Fisher Scientific). Lungs were removed en bloc and placed in a sterile 60 mm tissue culture dish (Catalog #CLS430166, Sigma-Aldrich) with an additional 2 ml of DMEM/F-12 + enzyme solution and cut into small pieces using sterile disposable surgical scalpels (Feather Safety Razor Co., Ltd, Osaka, Japan). Lungs were transferred to a 15 ml sterile tube, vortexed at medium speed periodically to help dissociate cells, and kept in a 37°C water bath for 30 minutes. Enzymes were inactivated by adding 5 ml DMEM/F-12 + 10% HyClone fetal bovine serum (Catalog #SH3091003, Fisher Scientific). Dissociated primary cells were spun down at 1000 rpm for 5 minutes, washed twice with PBS.

Samples were then strained through a 40 µm Corning Sterile Cell Strainer to obtain a single-cell suspension, and then incubated in 3% paraformaldehyde (Cat #: 28906, Thermo Fisher Scientific) for 30 minutes. Fixed and dissociated lung cells were strained again through a 40 µm sterile cell strainer, washed with PBS containing 0.1% Triton-X-100, counted, and 10 million cells were incubated with 10 µg of FITC-conjugated γ-H2AX(Ser139) primary antibody (yH2AX-FITC, EMD Millipore, Burlington, MA) on ice for 30 minutes. Cells were spun down and the cell pellet was washed twice with PBS containing 0.1% Triton-X-100, followed by final suspension in PBS containing 2% FBS. DRAQ5 nuclear stain (Thermo Fisher) was added at 5 uM 5 minutes prior to placement of samples in an ImageStreamX imaging flow cytometer (Amnis, Seattle, WA). Quantitative analysis and imaging of γ-H2AX foci (FITC: 570-620nm) and nuclei (DRAQ5: >655 nm) was performed on 10,000 cells per mouse, with bright field and fluorescence images quantified in separate channels and then merged. A compensation matrix was established with γ-H2AX and DRAQ5 stains in separate samples to eliminate overlap in emission spectra between channels. The IDEAS Spot Count analysis program was used to quantify γ-H2AX foci and γ-H2AX channel fluorescence. DRAQ5 fluorescence was used to ensure that all spots counted were in the nucleus.

### 2.6. The 8-isoprostane ROS assay

8-isoprostane was measured as an indicator of ROS in mouse lungs using an 8-isoprostane ELISA kit (Catalog # ab175819, Abcam). All mouse treatments were performed in triplicate and ELISA was performed in duplicate. Additional reagents used: 2N Sulfuric Acid stop solution (catalog # DY994, R&D Systems, Minneapolis, MN). Triphenylphosphine (TPP) (Catalog # T84409-1G, Sigma-Aldrich), and ethyl acetate (Catalog # 270989, Sigma-Aldrich). WT, *Alkbh8^def^* and *Cyp2abfgs-null/Alkbh8^def^* double mutant mice were sacrificed via CO_2_ asphyxiation at basal conditions and after challenge with 200mg/kg NA or corn oil carrier for 24h. Lungs were excised and divided among 2 tubes with each containing ½ a lung, flash frozen and stored at -80°C. The right lobes (on the right as observed with the mouse ventral-side up) were processed for 8-isoprostane measurements as outlined in the manufacturer protocol. A few modifications were made to the protocol to adjust for low sample volume, as described below. The left lobes were processed for glutathione quantification (Section 2.7).

Lungs were carefully weighed and then homogenized in 500uL of H_2_O containing 0.00125mg TPP (instead of 4 mL H_2_O and 0.01mg TPP recommended in the manufacturer protocol, due to smaller amount of available sample). 1ul of glacial acetic acid was added to acidify the samples, and pH measured using Hydrion S/R pH test paper (Micro Essential Labs, Inc., Brooklyn, NY) to ensure pH between 3-4. 500 uL ethyl acetate was added, sample vortexed, spun down at 5000 rpm for 5 minutes, and organic phase separated into a new tube. The ethyl acetate extraction was repeated 2 more times and organic phases pooled. The pooled organic phases were carefully dried using nitrogen gas. 5uL of ethanol, absolute (200 Proof) (Catalog #BP2818100, Fisher Scientific) was added to each sample to dissolve the dried residue. 300 uL 1X Sample Dilution Buffer (provided in the kit) was added to each sample. Samples were spun at 10,000 rpm for 5 min, and the supernatant removed and placed in a new 1.5ml microcentrifuge tube. The supernatant was used for ELISA directly following kit instructions.

### 2.7. Glutathione quantification

#### 2.7.1 Sample preparation

WT and *Alkbh8^def^*, N = 3, untreated and 24h corn oil or NA treated were sacrificed via CO_2_ asphyxiation and the lungs removed and immediately flash frozen. Samples were stored on dry ice and GSH & GSSG quantification was determined using liquid chromatography-tandem mass spectrometry (LC-MS/MS). Tissue samples were homogenized in 19x volumes of Tris-Acetate buffer (100mM Tris-base, 1.0mM ethylenediaminetetraacetic acid (EDTA), 150mM potassium chloride (KCl), pH 7.4) using a Polytron PT 10-35-GT powered homogenizer (Kinematica, Bohemia, NY).

Tissue homogenates were processed using protein precipitation and centrifugation as follows: 40µL internal standard, GSH-^13^C2,^15^N (3 µg/mL in 10% acetonitrile), was added to 40µL of tissue homogenate, followed by the addition of 200µL methanol and 200µL water. The samples were mixed and centrifuged at 4°C and 14,000 rpm for 10 minutes. The supernatant was collected. 300µL 0.1N HCL was added to 100µL supernatant and vortexed for 30 s and centrifuged at 4°C and 14,000 rpm for 10 minutes. The supernatant was collected and 3 μL injected into the LC-MS system.

#### 2.7.2 LC-MS conditions for analysis of GSH and GSSG

The LC-MS system consisted of an Agilent HPLC-1290 (Agilent, USA), and a Qtrap 6500 plus mass spectrometer (Sciex, Ontario, Canada) equipped with a Turbo IonSpray source. For data acquisition and processing, Analyst Software 1.6.3 was used.

Chromatographic separation was performed on a reversed-phase ZORBAX SB-C8 column (2.1 × 100 mm, 1.8µm) using a mobile phase consisted of 0.1% formic acid(A) and methanol(B) at a flow rate of 0.25 mL/min. The following mobile phase gradient was used: at 1%B, 0-3.5 min; 1%-50%B change, 3.5-4.0 min; 50%-95%B change, 4.0-4.5 min; at 95%B, 4.5-6.0 min; 95%-1%B change, 6.0-6.1 min; re-equilibration of the column at 1%, 6.1-9.5 min. The mass spectrometer was operated in positive ionization mode with an electrospray ionization probe and using multiple reaction monitoring (MRM). The MRM transitions used for detection of GSH and GSSG were 308.0/179.0 and 613.1/355.1, respectively. Isotopically labeled GSH (GSH-^13^C2,^15^N) was detected with an MRM transition of 311.0/182.0.

#### 2.7.3 Calibration curves

Calibration standards for GSH and GSSG were prepared by diluting the stock solutions with water:acetonitrile (9:1,v/v) to the final concentrations of 1, 2, 10, 20, 50, 100 µg/mL;. Calibration standards were processed using protein precipitation and centrifugation as follows: 40µL internal standard, GSH-^13^C2,^15^N (3 µg/mL in 10% acetonitrile), was added to 40µL of calibration standards and followed by the addition of 200µL methanol and 200µL water. The samples were mixed and centrifuged at 4°C and 14,000 rpm for 10 minutes. The supernatant was collected. 300µL 0.1N HCL was added to 100µL supernatant and vortexed for 30 s and centrifuged at 4°C and 14,000 rpm for 10 minutes. The supernatant was collected and 3 μL injected into the LC-MS system.

### 2.8. Protein assays

A Simple Western (Wes) protein analysis tool was used to quantify protein levels following manufacturer protocols (Protein Simple, Kanata, ON, Canada). Commercial antibodies in this study were as follows: GPX1 (Catalog #AF3798, R&D Systems, Minneapolis, MN), GPX3 (Catalog #AF4199, R&D Systems), TRXR1 (Catalog #MAB7428), TRXR2 (Catalog #ab180493, Abcam), TRX2 (Catalog #MAB5765), SELS (Catalog #HPA010025, Sigma) and GAPDH (Catalog # CB1001, Millipore Sigma). Lungs were homogenized in 1-1.5ml RIPA Buffer (50 mM Tris-HCl, pH 7.4, with 150 mM NaCl, 1% TridonX-100, 0.5% sodium deoxycholate, and 0.1% sodium dodecyl sulfate) supplemented with an EDTA-free phosphatase inhibitor cocktail 2 (Catalog #P5726, Sigma) and phosphatase inhibitor cocktail 3 (Catalog #P0044, Sigma). Extracted protein was quantified using either Bradford assay (BioRad, Portland, OR) or Pierce BCA Protein Assay Kit (Thermo Fisher) according to manufacturer’s instructions, then diluted to equal concentrations and, due to Wes’ sensitivity, measured a second time using the BCA assay, and adjusted if necessary. The final loading concentration of lysate into each capillary was 3 ul of 1.5 mg/ml lung protein.

### 2.9 Bronchoalveolar Lavage

WT, *Alkbh8^def^* and *Cyp2abfgs-null/Alkbh8^def^* were euthanized using CO_2_ asphyxiation, the lungs and trachea were exposed, a small incision was placed in the trachea, intubated using a sterile, disposable blunt end needle, 22 gauge, 0.5 in (Part #B22-50, SAI Infusion Technologies, Lake Villa, IL) secured with a piece of thread. Next, the lungs were slowly inflated with 0.8 ml PBS, using a 1-ml Coviden™ Monoject™ tuberculin syringe (Fisher Scientific, Franklin, MA) and fluid removed and re-injected 2 additional times over the course of ∼1 minute to draw out the maximum number of inflammatory cells. The PBS containing cells was then transferred to a 2-ml microcentrifuge tube. The PBS lavage step was repeated 2 more times with 0.5ml PBS each. The pooled Bronchoalveolar Lavage Fluid (BALF) was then mixed by gentle pipetting, dissolved 1:1 with trypan blue (Thermo Fisher), and live cells immediately counted on a hemocytometer. Cells/ml were plotted (N = 3 mice per genotype per condition), with error bars denoting standard error of the mean.

## Results

### Alkbh8^def^ lungs have adapted by molecular reprogramming

Under standard care conditions, with no external applied stress, 12-week-old WT and *Alkbh8^def^* mice (Fig. 2A) and lungs have no observable differences (Fig.2B). Similarly, H&E stained lung sections of the bronchioles and alveolar regions do not show any noticeable difference (Fig. 2C). Lung sections from WT and *Alkbh8^def^* mice show healthy epithelial cells, comprised primarily of club cells and ciliated cells, lining the bronchioles. Epithelial cells maintain a cuboidal structure and line the perimeter of the bronchiole, with club cells distinguishable from ciliated cells by the absence of cilia on the surface facing the bronchiolar lumen. No inflammation (i.e. macrophages present) or swelling in the alveolar regions was observed in either genotype. Together, these results support that even though there is a deficiency in *Alkbh8* (Fig. 2D) mouse development and lung physiology appear normal under normal growth conditions.

**Figure 2.**
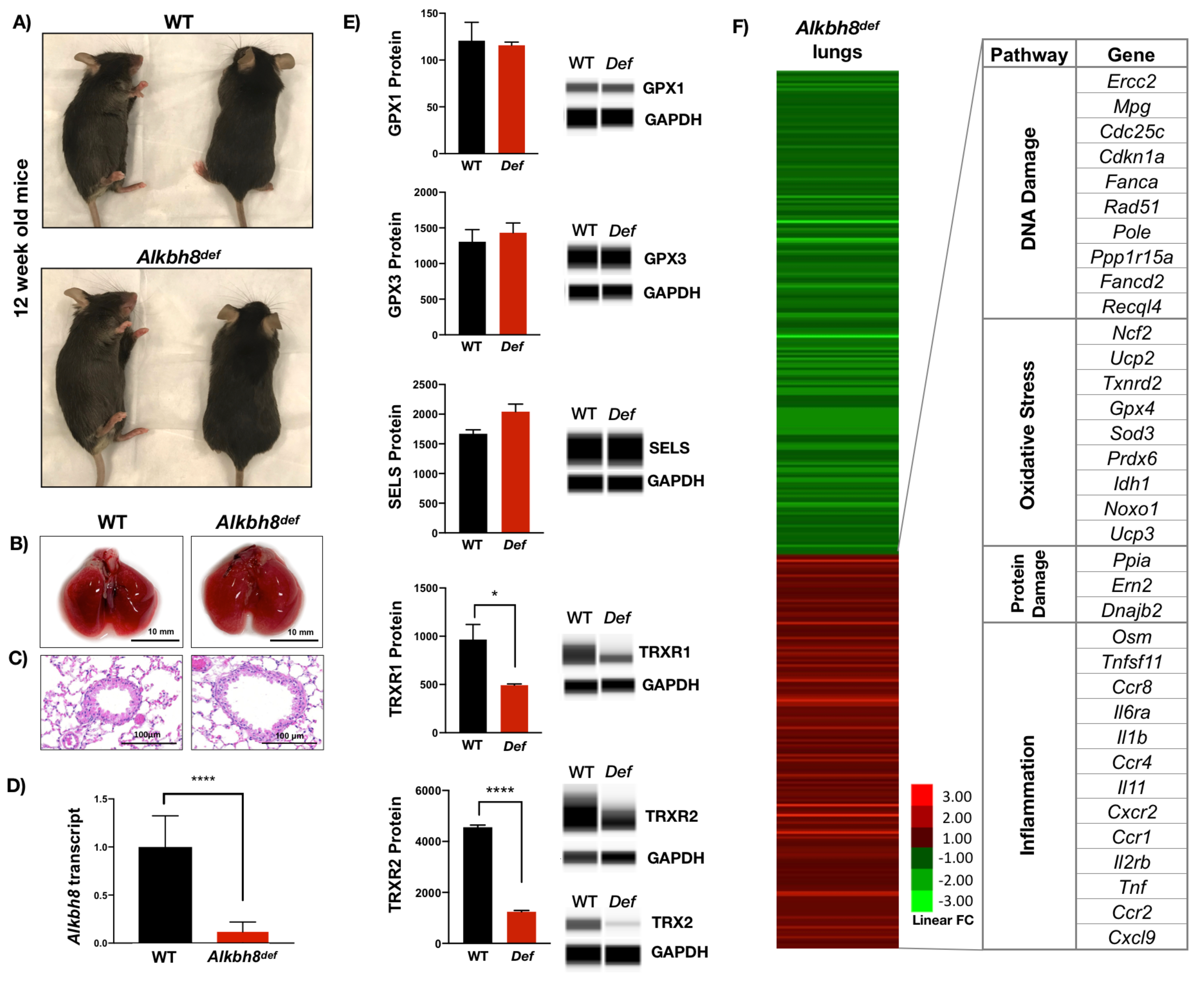
*Alkbh8^def^* lungs have decreased levels of specific selenoproteins and transcriptionally reprogram stress response systems. **A)** 12 week old mice, **B)** lungs and **C)** hematoxylin and eosin stained lung sections, showing bronchioles and alveolar regions, from WT and *Alkbh8^de^*^f^ mice. **D)** *Alkbh8* transcript levels from basal WT and *Alkbh8^def^* lungs were quantified using qPCR analysis (N = 3 per genotype, P ≤ 0.0001). **E)** GPX1, GPX3, SELS, TRXR1 and TRXR2 selenoprotein expression levels and non-selenoprotein TRX2 (blot only) from basal WT and *Alkbh8^def^* lungs were assed using Wes® Simple Western. **F)** Targeted RNA sequencing was performed on 350 stress response genes and fold differences in number of sequence reads after normalization to endogenous control (Ribosomal protein L13a, Rpl13a) was used to generate a heat map. Expression values are *Alkbh8^def^* lungs relative to WT under basal conditions. Notable transcripts with fold-change values of 1.5 or greater in each signaling pathway are listed. N = 2 mice per genotype.

We also investigated molecular changes in the lungs of *Alkbh8^def^* mice by measuring selenoprotein levels. We used a Simple Western (Wes) automated western blot tool for protein detection. Under normal growth conditions, we observed little difference in GPX1, GPX3, and Selenoprotein S (SELS) protein levels. In contrast we observed a significant decrease in TRXR1, TRXR2 and TRX2 protein levels in the lungs of *Alkbh8^def^* mice. TRXRs are Sec-containing proteins critical for all signaling pathways in which thioredoxin (TRX) is involved as a reducing agent as TRXRs maintain TRX in its reduced and active state. As such, they support other TRX-dependent antioxidants such as methionine sulfoxide reductases (MSRS) and peroxiredoxins (PRXs). Dysfunctions in the TRXR system have been linked to oxidative stress, mitochondrial dysfunction, apoptosis and sensitivity to added stress (Lopert, Day, & Patel, 2012), and molecular reprogramming in the lung to adapt to the stress (Locy et al., 2012). TRX2 is localized to the mitochondria and is an essential member of the thioredoxin system, with deletions associated with massive dysregulation of redox balance, apoptosis and early embryonic lethality in mice, corresponding to the time when mitochondria develop to begin oxidative phosphorylation(Nonn, Williams, Erickson, & Powis, 2003; Tanaka et al., 2002). TRX2 is a non-selenoprotein enzyme containing cysteine in the active site and functions to maintain protein thiols in a reduced state, thereby becoming oxidized. As such, we predicted that molecular reprogramming of ROS and protein-damage response genes may play a factor in *Alkbh8^de^*^f^ mouse lungs in response to TRX2 and TRXR1 and TRXR2 deficiency.

Next we used mRNA-seq (Fig. 2F) to determine if other stress response pathways are transcriptionally regulated to adapt to the *Alkbh8* deficiency and corresponding decrease in antioxidant capacity. Many transcripts belonging to inflammatory pathways were upregulated in the lungs of *Alkbh8^def^* mice, relative to WT. For example, oncostatin M (*Osm*) was up-regulated 6.82 fold in in the lungs of *Alkbh8^def^* mice. OSM is a secreted cytokine and functions to regulate other cytokines. As such, we also observed an increase in transcripts specific to *Ccr8* (2.5 fold), *Il6ra* (2.24 fold), *Il1b* (2 fold), *Ccr4* (1.82 fold), *Il11* (1.82 fold), *Cxcr2* (1.75 fold), *Ccr1* (1.67 fold), as well as others in the in the lungs of *Alkbh8^def^* mice. These proteins functionally participate in the inflammatory and immune response and include members of the G-protein-coupled cytokine receptor family (CCRs, CXCRs), the interleukin 6 receptor subunit (IL6RA), and inflammatory cytokines IL1B and IL11, both of which have pleiotropic effects supporting the immune system. In addition, we observed protein damage transcripts upregulated in *Alkbh8^def^* mice including *Ppia* (2.78 fold), *Ern2* (1.77 fold), and *Dnajb2* (1.75 fold) relative to WT. PPIAse proteins function in the cis-trans isomerization of peptide bonds and aid in protein folding, ERN2 participates in signaling between the nucleus and endoplasmic reticulum, and DNAJB2 is part of the heat shock protein family.

Transcripts specific to DNA damage and oxidative stress were also upregulated in the lungs of *Alkbh8^def^* mice, relative to WT. For example, the transcript corresponding to the nucleotide excision repair protein ERCC2 and base excision repair protein MPG were up-regulated 2.48 and 2.29 fold, respectively, in the lungs of *Alkbh8^def^*. Other DNA repair transcripts upregulated in the lungs of *Alkbh8^def^* mice include those for *Rad51* (1.75 fold), *Pole* (1.7 fold), and *Fancd2* (1.5 fold). In the oxidative stress pathway, the highest increase in *Alkbh8^def^* mice was in Neutrophil Cytosolic Factor 2 (*Ncf2,* 1.79 fold), a subunit of a multi-protein NADPH oxidase complex necessary for superoxide production in cells. Mitochondrial Uncoupling Protein 2 (*Ucp2*, 1.71 fold) was also increased in the lungs of *Alkbh8^def^* mice. UCP2 functions in separating oxidative phosphorylation from ATP synthesis with energy released as heat and it plays a role in thermogenesis, diabetes and obesity. The *Trxr2* (1.69 fold) and *Gpx4* (1.54 fold) transcripts, which encode ROS detoxifying selenoproteins as well as Superoxide Dismutase 3 (*Sod3*, 1.53 fold), were also upregulated. As selenoproteins are translationally regulated by ALKBH8, the increase in transcript levels may be a compensatory mechanism in response to decreased protein levels. Together the increased levels of transcripts specific to DNA repair and oxidative stress supports the idea that the lungs of *Alkbh8^def^* mice have an increased stress and DNA damage.

Oxidation-reduction potential (ORP) is a direct measure of the balance between ROS and antioxidants in a system. The more positive the ORP, the higher the oxidant activity or alternately, the lower the antioxidant activity. ORP is considered a new marker of ROS and is used clinically as a measure of oxidative stress in blood of patients with traumatic brain injury, stroke, sepsis, metabolic disorder and liver toxicity (Bar-Or, 2009; Bjugstad & Fanale, 2016; Bjugstad et al., 2016; Bobe et al., 2017; Rael et al., 2007; Spanidis et al., 2015). Isoprostanes are a family of eicosanoids produced by the random oxidation of tissue phospholipids by ROS and provide one of the most reliable measures of oxidative stress in vivo (Gross et al., 2005; Morrow, 2005). We observed approximately 10% increase in oxidation reduction potential (ORP) (Fig. 3A) and a near 40% increase in the average 8-isoprostane levels (Fig. 3B) in the lungs of *Alkbh8^def^* mice. We did not observe differences in 2’,7’-dichlorofluorescin diacetate (DCFDA) in *Alkbh8^def^* mice (Supplemental Fig. 2A-B), but this is a less sensitive assay compared to 8-isoprostane levels. Lastly, we measured GSH and GSSG in basal mouse lungs and plotted the ratio as an indicator of oxidative stress. The *Alkbh8^def^* mice have less than half of the GSH/GSSG ratio under basal conditions (Fig. 3C). Separate graphs for GSH, GSSG and total glutathione can be found in Supplemental Figure 3. Together, our ORP, 8-isoprostane and GSH/GSSG findings support the idea that there is increased ROS in the lungs of *Alkbh8^def^* mice and possibly a GSH synthesis defect.

**Figure 3.**
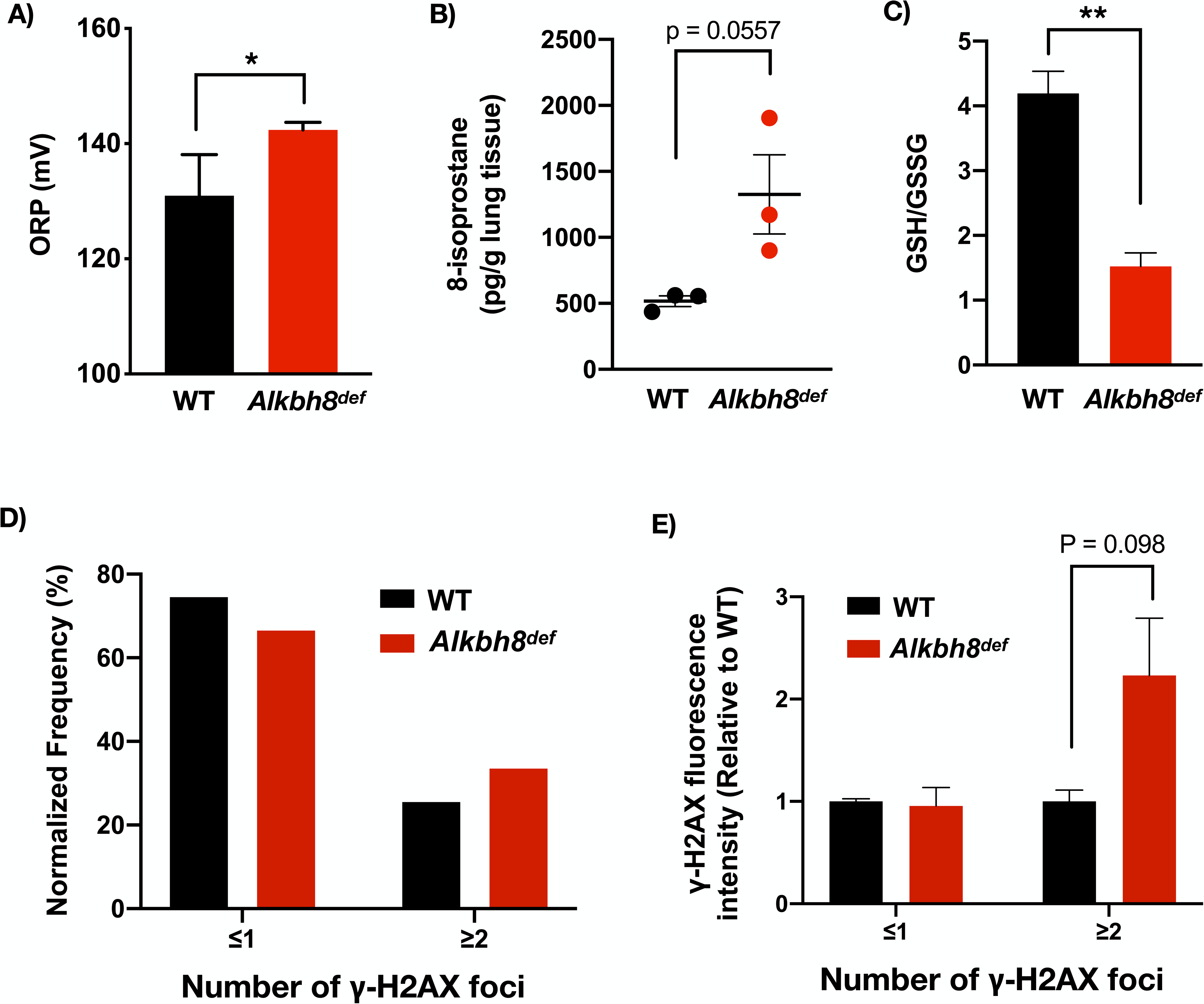
Markers of oxidative stress and increased DNA damage response observed in *Alkbh8^def^* lungs. **A)** Oxidation reduction measurements in the form of ORP were performed on WT and *Alkbh8^def^* lungs and we observed a significant difference (P ≤ 0.05, N = 3). **B)** 8-isoprostane levels (pg/gram lung tissue) were increased in *Alkbh8^def^* lungs (P = 0.0557, N = 3). **C)** GSH and GSSG were measured using LC-MS/MS and plotted as a ratio, with *Alkbh8^def^* mice showing a decrease (N = 3, P ≤ 0.01) **D)** Basal WT and *Alkbh8^def^* lungs were excised and enzymatically and mechanically separated into a single-cell suspension, fixed and stained with a FITC conjugated antibody specific for γ-H2AX and the DRAQ5 nuclear stain, followed by quantitative imaging with an Amnis Imagestream ISX100 flow cytometer. γ-H2AX foci were scored for intensity and number using the IDEAS Spot Count analysis program, presented as the frequency of cells with ≤1 spots or ≥2 spots (N = 3 mice per genotype, 10,000 cells per lung). **E)** γ-H2AX-FITC channel fluorescence intensity was quantified and normalized to WT to demonstrate there are increased levels in *Alkbh8^def^* lungs (P = 0.098).

Increased ROS can promote increased DNA strand breaks (Møller & Wallin, 1998; Sharma et al., 2016). γ-H2AX is recruited to sites of double stranded DNA breaks by ATM and is a classic marker for the DNA damage response (DDR) (Rogakou, Pilch, Orr, Ivanova, & Bonner, 1998). We assayed for γ-H2AX using dissociated primary lung cells processed through an Amnis Imagestream imaging flow cytometer. The Amnis Imagestream takes single-cell brightfield and fluorescent images, which can be spatially quantitated and set to count the number of foci in a cell. We processed 10,000 dissociated primary lung cells assayed with a γ-H2AX antibody. Example brightfield and fluorescent images are shown in Supplemental Fig. 2C for low γ-H2AX spot count and 2D for high spot count. We observed an increase in γ-H2AX foci in the lungs of *Alkbh8^def^* mice, relative to WT. The number of *Alkbh8^def^* lung cells show a shift toward increased spot counts (Fig. 3D), as well as increased γ-H2AX channel fluorescence in the ≥2 spot count group (Fig. 3E), supporting our earlier finding of increased DNA damage response transcripts in *Alkbh8^def^* mice. Ultimately our transcript, ROS and DDR data support the idea that under normal growth conditions the lungs of *Alkbh8^def^* are experiencing some stress but have undergone molecular reprograming to adapt to the epitranscriptomic and selenoprotein deficiency. Importantly, the *Alkbh8^def^* mice are adapting to the stress by upregulating a variety of signaling pathways.

### Alkbh8^def^ mice show hypersensitivity to NA and have increased lung airway inflammation after NA exposure

We next asked whether the *Alkbh8^def^* mice would be sensitive to a challenge by the environmental toxicant NA. We treated WT, *Alkbh8^def^*, *Cyp2abfgs-null*, and *DM* (*Cyp2abfgs-null/Alkbh8^def^*) mice, the last two of which lack the CYPs required for NA bioactivation, to an acute 200 mg/kg dose of NA (Fig. 4). The 200 mg/kg dose via I.P. injection is well established in literature as the amount of NA delivered to the lung is relevant to OSHA standard for NA exposure in the workplace (10 ppm, or 15 ppm short-term). We observed overall appearance, lung morphology and quantified cells in the bronchoalveolar lavage fluid (BALF). *Alkbh8^def^* mice have an observable difference in appearance after NA exposure compared to WT or DM mice. They display more fluffed fur, which is a classic marker of stress in mice. Note that *DM* mice, despite *Alkbh8* deficiency, have a smooth coat of fur, supporting previous studies that NA bioactivation is required for toxicity (Buckpitt & Warren, 1983; Warren, Brown, & Buckpitt, 1982). Figure 4C shows representative lung airway images of all four strains of mice, either untreated or treated with 24h corn oil or NA dose (200 mg/kg), with H&E stained lung sections focused on the bronchioles (round structures) and alveolar regions (perimeter). Healthy lung airways were observed under untreated and corn oil conditions in all 4 strains. After NA exposure, the *Cyp2abfgs-null* and *DM* mice are resistant to NA-induced pulmonary damage. WT mice show a flattening and elongating of epithelial cells, an event that occurs after early NA-induced necrosis of club cells (Plopper et al., 2013a). Club cells participate in the biotransformation of many harmful substances introduced into the lung through inhalation, including NA, via CYP-mediated bioactivation. The lung airway of *Alkbh8^def^* mice appear altered relative to WT, as what appears to be necrotic cells were observed in the airway epithelium. Lungs were sectioned, H&E stained, and analyzed by a board-certified pathologist as described in the methods. Pathologies associated with NA-induced lung include free floating necrotic cells in the airway lumen, flattened and elongated club cells in the airway epithelium, absent club cells in parts of the airway, and increased presence of macrophages. The pathologies were more extreme in *Alkbh8^def^* mice, which was further increased at prolonged exposure durations, discussed in further detail in a later section. An acute 24-hour dose of NA (200mg/kg) resulted in elongated club cells, a classic response to NA, in most WT and *Alkbh8^def^* mice. However, the *Alkbh8^def^* mice exhibit necrosis of epithelial cells in many more bronchioles. We next used bronchoalveolar lavage to measure the number of cells in the recovered fluid, so as to gauge inflammation in WT, *Alkbh8^def^* and *DM* mice (Fig. 4D). *Alkbh8^def^* mice have a significantly increased number of cells in the bronchoalveolar lavage fluid (BALF) compared to WT mice, under both corn oil and NA treated conditions (P < 0.01, P < 0.05, respectively).

**Figure 4.**
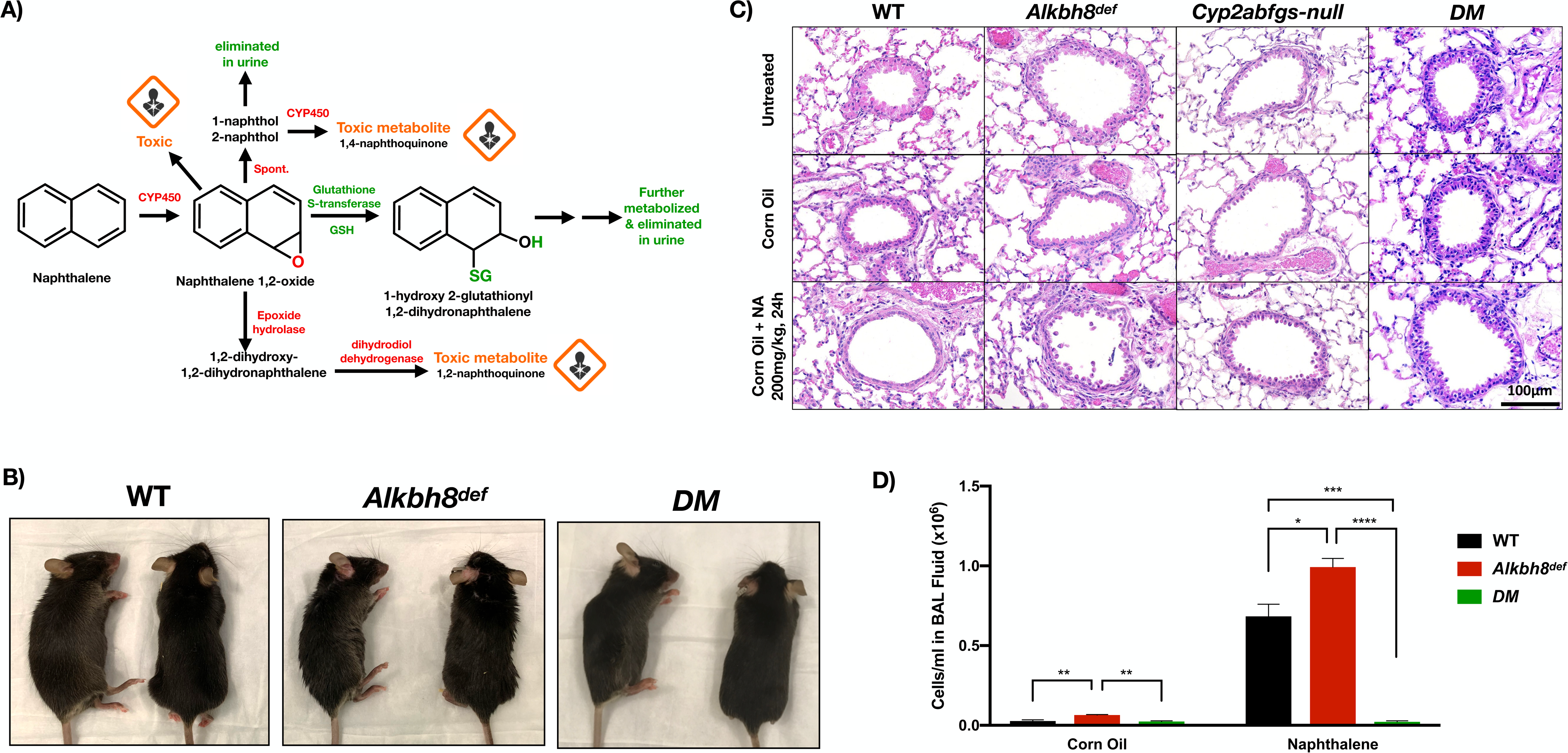
*Alkbh8^def^* mice and lungs are more sensitive to damage from NA. **A)** NA is bioactivated by CYP enzymes into naphthalene 1,2-oxide, a reactive epoxide metabolite, which undergoes further metabolism. **B)** Representative images of mice treated with NA (200 mg/kg) for 2 days. **C)** Hematoxylin and eosin stained lung sections showing bronchioles and alveolar regions from WT, *Alkbh8^de^*^f^, *Cyp2abfgs-null*, and *Alkbh8^def^/Cyp2abfgs*-null double mutant (DM) mice under untreated (first row), 24h corn oil carrier (second row), and 24h NA 200mg/kg (third row) conditions (N = 3 per genotype & condition). **D)** The concentration of cells (cells/ml) in bronchoalveolar lavage fluid obtained after 24h corn oil or NA treatment (200 mg/kg). (N = 3 mice per condition, error bars denote SEM. * P ≤ 0.05, ** P ≤ 0.01, *** P ≤ 0.001, **** p ≤ 0.0001.)

### Alkbh8^def^ mice show an increase in molecular markers of stress compared to WT mice under NA challenged conditions

Next we investigated the redox and transcriptional states of WT, *Alkbh8^def^* and DM mice 24 hours after NA. ORP measures highlight a NA-induced increase in oxidants, or a deficiency in antioxidants, in *Alkbh8^def^* mice (P = 0.058), while a reduction is observed in WT mice (P ≤ 0.05) (Fig. 5A). We note that the increase in ORP observed in Fig. 3A in untreated *Alkbh8^def^* mice is no longer seen in the corn oil condition. Although relative values are presented, we attribute this to a slight increase in WT ORP after corn oil injection while *Alkbh8^def^* ORP remained the same (raw data not shown). This may suggest some metabolic differences between WT and *Alkbh8^def^* mice. While ORP is a general indicator of oxidative stress, a more specific *in vivo* measure is 8-isoprostane levels. We observed an increase in 8-isoprostane levels in both WT and *Alkbh8^def^* mice treated with NA, relative to the untreated sample (Fig. 5B). WT NA-treated mice show a statistically significant increase (P ≤ 0.05) in 8-isoprostane levels relative to the corn-oil treated WT group, while NA-treated *Alkbh8^def^* mice show a more significant increase (P ≤ 0.001) relative to corn oil-treated mice. Although *Alkbh8^def^* mice show an increase in 8-isoprostanes in the lung, the difference relative to the WT NA-challenged mice is not statistically significant (P = 0.23). *DM* mice do not have notable differences in 8-isoprostane levels in the lung between corn oil and NA treatment conditions. We again note a small but statistically significant difference in the corn oil condition between WT and *Alkbh8^def^*, supporting the idea of metabolic differences between the mice. We further measured the GSH and GSSG in CO and 200mg/kg NA-treated mouse lungs after 24 hour exposure. We observe a decreased GSH/GSSG ratio in *Alkbh8^def^* mice under both conditions (Fig. 5C), again supporting the idea of increased oxidative stress in the writer-deficient mice.

**Figure 5.**
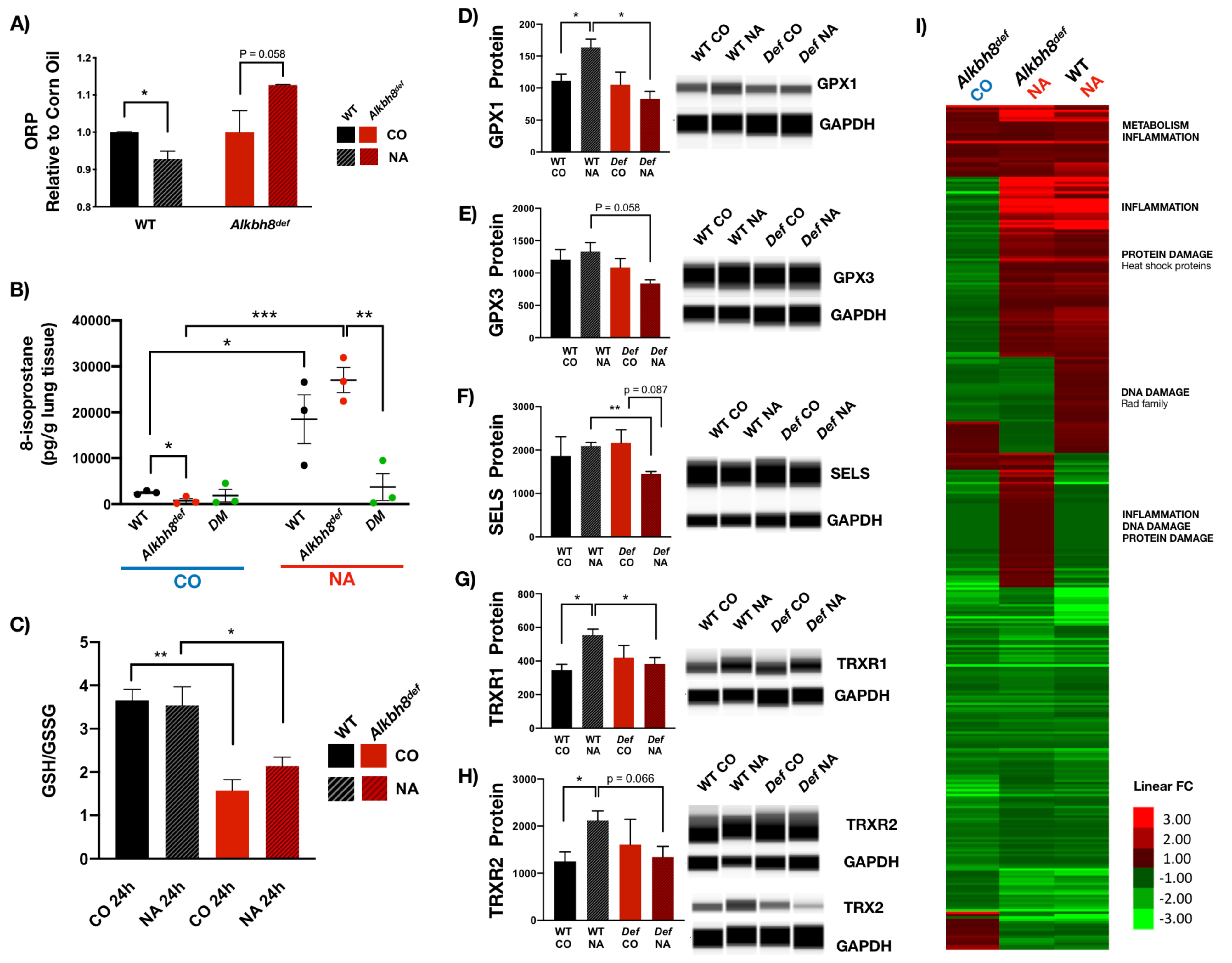
Altered stress response to NA observed in *Alkbh8^def^* lungs. **A)** ORP measurement in WT and *Alkbh8^def^* lungs 24 hour after NA exposure, 200mg/kg. (N = 3 mouse lungs per genotype and treatment condition) **B)** 8-isoprostane levels (pg/g lung tissue) are plotted as described in figure 3B. (N = 3, error bar denotes SEM. * P ≤ 0.05, ** P ≤ 0.01, *** P ≤ 0.001, N = 3 mouse lungs per condition) **C)** GSH and GSSG were measured using LC-MS/MS and plotted as a ratio, with *Alkbh8^def^* mice showing a decreased ratio under both CO and NA-treated conditions. (* P ≤ 0.05, ** P ≤ 0.01, N = 3 mouse lungs per condition.) **D)** Protein levels of GPX1 **E)** GPX3, **F)** SELS, **G)** TRXR1 and **H)** TRXR2 and TRX2 (blot only) were determined using the Wes system. GAPDH loading control was multiplexed with each antibody and is shown at the bottom in each representative blot. Both targets (selenoprotein & GAPDH) were multiplexed within the same capillary. (N = 3 mice per genotype. *P ≤ 0.05, ** P ≤ 0.01.) **I)** Heat map showing differences in quantitated gene expression of 350 stress response genes via targeted RNA sequencing. Comparison after 24h corn oil or 200mg/kg NA exposure, with some functional highlights outlined on the right. All comparisons are relative to WT corn oil treatment. A large version of the heat map is available in Supplementary Fig 1. (N = 2 per condition, per genotype).

We also measured protein levels to determine if specific selenoproteins are upregulated in response to NA and whether this was corrupted in the *Alkbh8^def^* lungs. WT mice upregulate the selenoproteins GPX1, TRXR1 and TRXR2, and non-selenoprotein TRX2 in response to NA stress (Fig. 5D-H). In contrast we observed a consistent failure to upregulate selenoprotein levels in *Alkbh8^def^* mice (Fig. 5D-H, with full blots available in Supplemental Figure 5). While we note decreased TRXR1 and 2 in *Alkbh8^def^* lungs under basal conditions (Fig. 2E), after corn oil treatment the levels are similar to WT corn oil treated lungs again suggesting that there may be some metabolic effect or some level of oxidants present in the oil causing the *Alkbh8^def^* to upregulate TRXR proteins from baseline. Together the data support a model in which there are increased ROS levels and decreased ROS detoxifiers in the *Alkbh8^def^* lungs.

We also used targeted mRNA sequencing of mouse lungs to determine if the transcriptional response to NA was altered in the *Alkbh8^def^*mice, relative to WT. We observed distinct groups of transcripts that were up-regulated in WT but down-regulated in *Alkbh8^def^*, and vice versa (Fig. 5I **&** Table 1). Inflammatory markers were greatly increased in the lungs of *Alkbh8^def^* mice, relative to WT, but other signaling pathways were also affected. DNA damage response marker Exonuclease 1 (*Exo1*) was increased in *Alkbh8^def^* mice relative to WT. Heat shock transcripts including *Hspa1α*, *Hspa1b*, and *Hsph1* were upregulated in the *Alkbh8^def^* mice, relative to WT, which suggests that there may be some protein damage occurring. Induction of pulmonary heat shock proteins after NA exposure has previously been demonstrated and is predicted to be a response to NA-protein adduction (Williams, Cruikshank, & Plopper, 2003).

**Table 1a.**
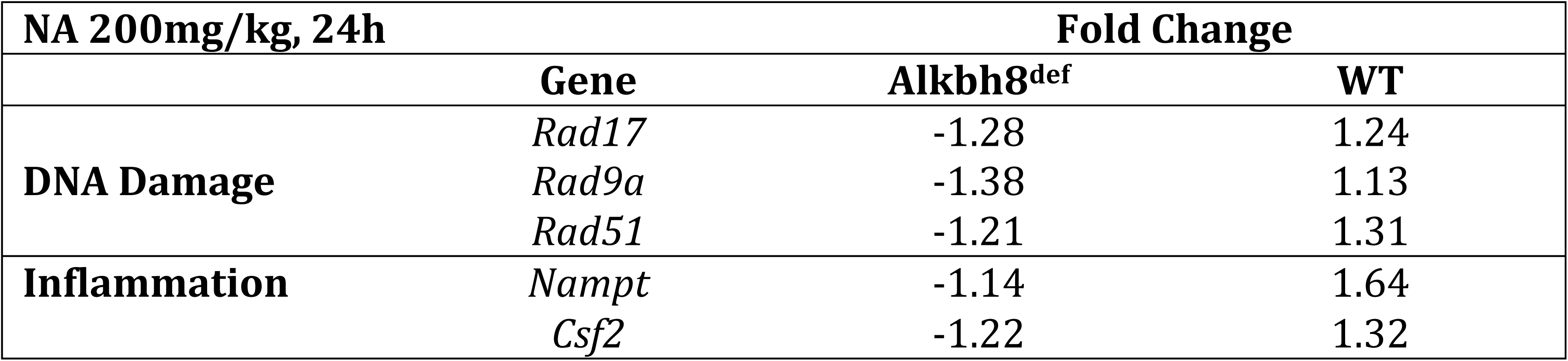
Transcripts upregulated in *Alkbh8^de^*^f^ mice but downregulated in WT in response to 24h NA exposure at 200mg/kg. Differences in response are for transcripts with greater than 0.5 fold difference between the two genotypes (N = 2).

**Table 1b.**
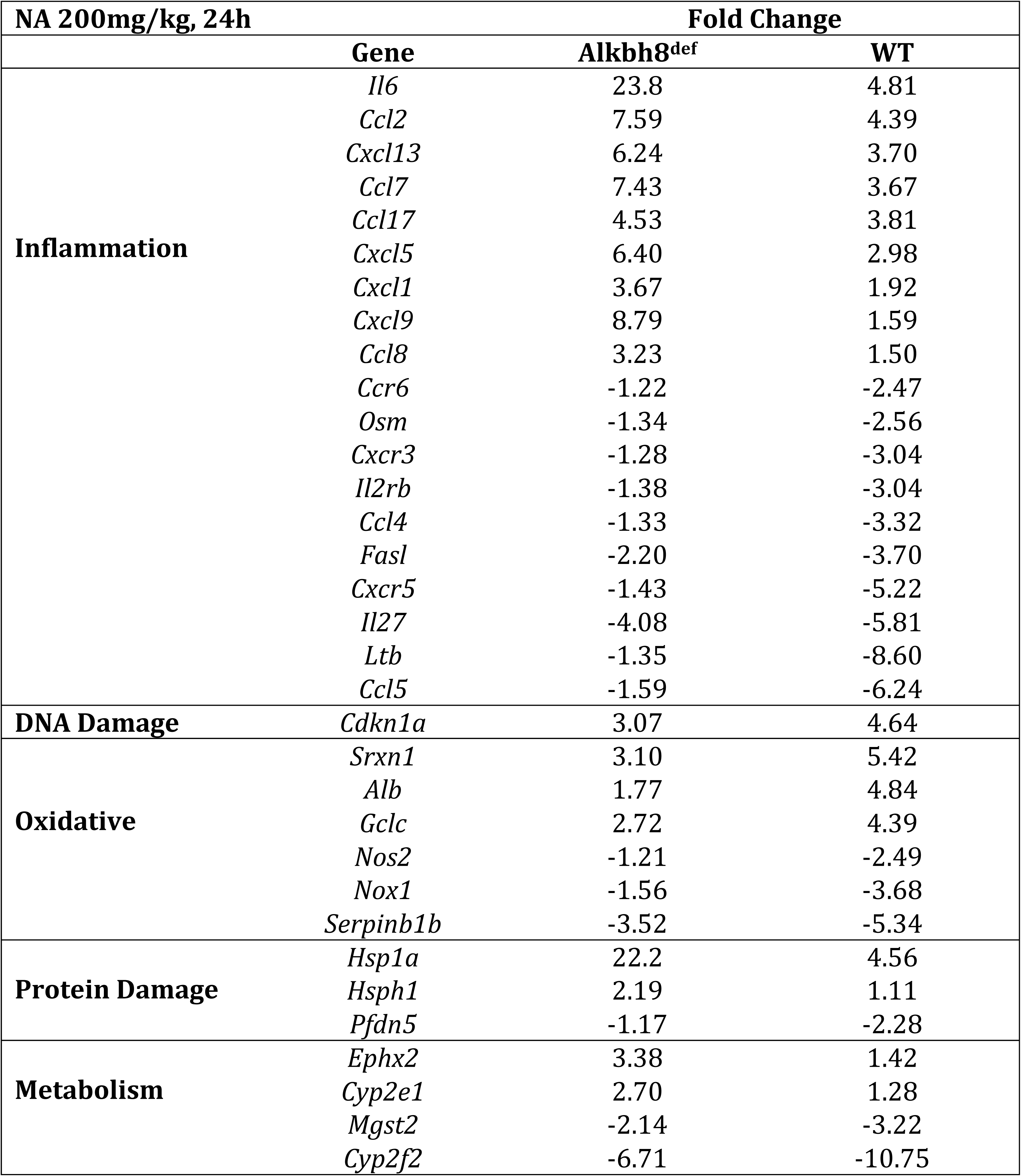
Transcripts downregulated in *Alkbh8^def^* mice but upregulated in WT in response to 24h NA exposure at 200mg/kg. Differences in response for transcripts with greater than 0.5 fold difference between the two genotypes (N = 2).

**Table 1c.**
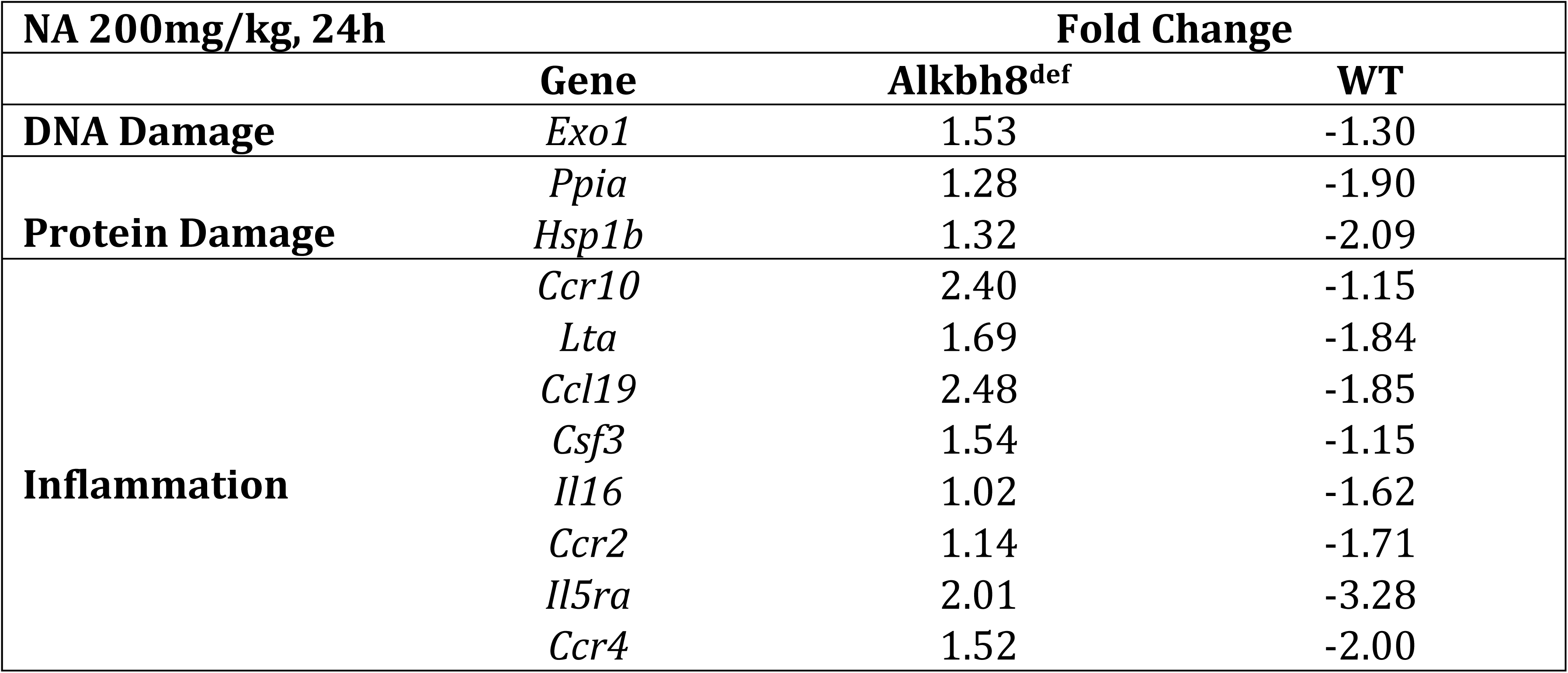
Transcripts with correlated responses (*Alkbh8^def^* and WT both up or both down) for transcripts with greater than 1-fold difference between the two genotypes (N=2).

Surprisingly, little difference in transcript levels for oxidative stress signaling pathway was observed when comparing the WT and the *Alkbh8^def^* mice at the acute dose. We predict that this is due to several reasons. First, *Alkbh8^def^* mice function under increased oxidative stress at basal conditions (Fig. 3A-C), which likely pre-conditions them for oxidative stress at acute doses due to molecular reprograming of oxidative stress response systems (Endres et al., 2015). Second, acute exposure to NA results in damage to the pulmonary club cells in the airway epithelium comprising only a small fraction of all lung cells. Due to the large cellular heterogeneity of the lung material used for RNAseq, this may lead to limited sensitivity of the assay to detect changes which are only occurring in club cells. As such, we note a much greater increase in inflammatory markers in *Alkbh8^def^* mice relative to WT mice in response to NA. As inflammatory cells, such as cytokine and chemokine-secreting macrophages, are recruited and distributed throughout the lung including airways and alveolar regions, RNAseq was able to detect the increases in inflammatory transcripts, which comprise by far the largest group of upregulated transcripts in response to NA treatment. For example, *Il6* had a 23.8 fold upregulation relative to WT mouse lungs. IL6 is produced at sites of tissue inflammation and is important in the response to acute lung injury (Gadient & Patterson, 1999; Voiriot et al., 2017). Other notable examples of cytokines and chemokines that have approximately double the increase in *Alkbh8^def^* mice relative to WT mice after NA exposure include *Ccl2, Ccl7, Ccl8, Ccl19, Cxcl1, Cxcl5, Cxcl9, Cxcl13, Ccr10*. Importantly, *Alkbh8* transcript remains decreased in *Alkbh8^def^* mice after CO and NA challenge (Supplemental Fig 4). These data support the idea that there is clear evidence of increased stress, inflammation and lung injury in the lungs of *Alkbh8^def^* mice 24 hours after NA.

### Alkbh8^def^ mice are sensitized to NA and fail to develop tolerance

We next used sub-chronic NA exposure conditions to determine whether there were differences in survival between WT and *Alkbh8^de^*^f^ mice. Mice were exposed to NA daily for two weeks to test their ability to survive and the resulting Kaplan-Meier plot (Fig. 6A) provides evidence that *Alkbh8^def^* mice have increased sensitivity to NA, relative to WT. We observed that over 50% of the WT mice survive and start gaining weight around day 3-4 with no deaths observed after day 5 (Fig. 6A-B). In contrast, all *Alkbh8^def^* mice die by day 7 and do not show marked recovery in weight. As the turnaround time for WT mice was around day 3, we sent lung tissues after 3-days of exposure to a board-certified pathologist for analysis. While WT mice had the early classic response of flattened and elongated club cells and some necrosis, they begin to recover the cuboidal structure of their airway epithelial cells around day 3, a sign of tolerance. Meanwhile, *Alkbh8^def^* lungs remain severely damaged with complete absence of epithelial cells in many parts of the bronchioles and injury and necrosis extending to ciliated cells to a moderate degree (Supplemental Fig. 6). The *Alkbh8^def^* mice also present blood clot formation in the alveoli suggesting either a greater incidence of hemolytic anemia, which NA is known to cause and which can contribute to intravascular coagulation, or injury to lung cells outside of the airways, such as endothelial cells in alveolar capillaries (ATSDR, 2005; Cappellini, 2007).

**Figure 6.**
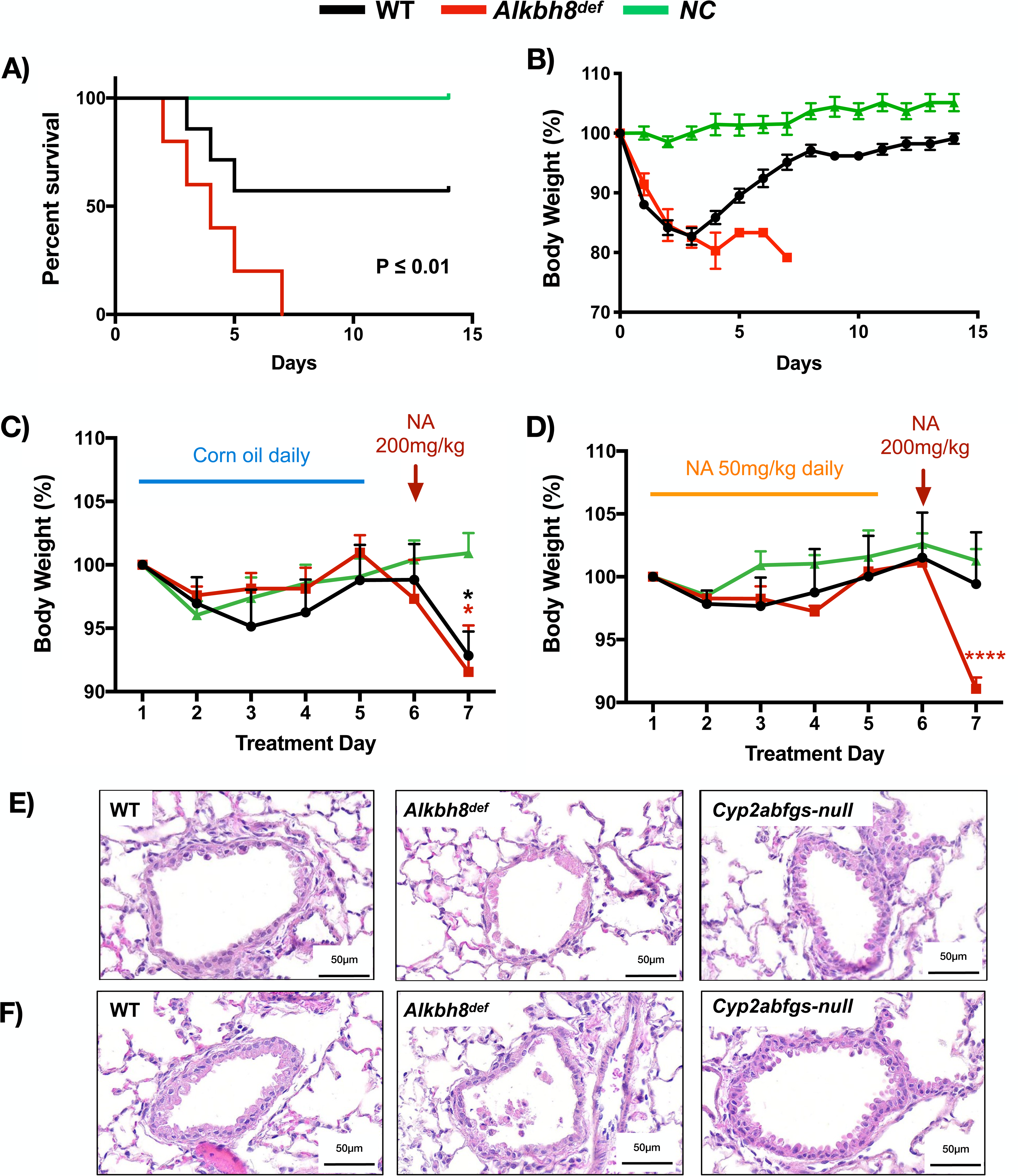
*Alkbh8* deficient mice show increased mortality and fail to develop tolerance to NA exposure. Mice were tested for their ability to tolerate a prolonged exposure to NA. **A)** Kaplan-Meier plot showing survival over 14 days after daily injection with 200 mg/kg NA. The log-rank (Mentel-Cox) test was used to generate a P-value. (P < 0.01). **B)** Weight was recorded daily over the 14-day study and plotted relative to starting weight. Any mice with a 20% or greater weight loss relative to starting weight were euthanized. **C)** Mice were tested for their ability to build tolerance to NA. The non adapted mice were treated with 5 doses of the carrier control, corn oil, followed by a 200 mg/kg high dose of NA. Average weight was plotted relative to starting weight. **D)** Mice were adapted to NA to build tolerance by subjecting them to 5 days of low-dose NA (50 mg/kg) followed by a 6^th^ day of high dose (200 mg/kg). Average weight was plotted relative to starting weight. Representative haemotoxylin & eosin (H&E) stained lungs shown for **E)** non-tolerated and **F)** NA tolerated WT, *Alkbh8^def^* and *DM* mice. Pictures show representative images of lung sections depicting bronchioles and alveolar regions (N = 3 for all genotypes and test conditions).

Based on these data WT mice appear to be developing tolerance to NA by day 3, and we performed dedicated tolerance study to investigate. As a control (non-tolerant) experiment we treated mice with 5 daily doses of corn oil, followed by a NA dose of 200 mg/kg. Both WT and *Alkbh8^def^* mice lost significant amounts of weight and show lung airway damage 24 hours after NA treatment (Fig. 6C&E), with WT and *Alkbh8^def^* lung airways showing a similar response to a single 24-hour NA treatment, as expected (Fig. 4C). *Cyp2abfgs-null* mice show no loss in weight after the NA dose, and show a small dip in weight early on after corn oil which we attribute to handling or injection stress. Next, we tolerated the mice with 5 daily doses of 50 mg/kg NA, followed by a high dose of 200 mg/kg on day 6 (Fig. 6D). As opposed to non-tolerated mice (5 corn oil treatments followed by a high dose of NA), NA tolerated WT mice did not lose significant weight. In contrast *Alkbh8^def^* mice lost a similar amount of weight as in the non-tolerant study. H&E stained lung sections from the tolerated mice provide evidence that the *Alkbh8^def^* mice have a marked increase in lung airway damage compared to WT and *Cyp2abfgs-null* mice, with continued necrosis and exfoliation of epithelial cells to the airway lumen in most bronchioles compared to WT mice (Fig. 6F).

## Discussion

We have previously shown that under normal growth conditions mouse embryonic fibroblasts deficient in the epitranscriptomic writer ALKBH8 have increased ROS and DNA damage but adapt to these stresses by molecular reprogramming stress-response transcript levels (Endres et al., 2015). Our current study supports the idea that this reprogramming can be extended to the lungs of *Alkbh8^def^* mice, as they have adapted to the epitranscriptomic deficiency and corresponding decrease in selenoprotein levels by upregulating the expression of inflammation, DNA repair and protein stress-response transcripts and markers. Molecular reprogramming of stress response systems in response to an epitranscriptomic writer deficiency has also been reported in lower organisms. For example, deficiencies in wobble uridine modification systems in budding yeast have been linked to upregulation of heat shock and unfolded protein response pathways (Patil et al., 2012). Molecular reprogramming due to cellular stress, including oxidative stress, protein stress and DNA damage, is also a well-documented response and has implications in development, aging and disease susceptibility. For example, the nuclear factor erythroid 2-related factor 2 (NRF2) is a master regulator of cellular resistance to oxidative stress(Qiang Ma, 2013; Vomund, Schäfer, Parnham, Brüne, & Von Knethen, 2017). In response to oxidative stress, NRF2 dissociates from its negative regulator KEAP1, translocates to the nucleus, and binds to specific antioxidant response elements (AREs) in the promoter region of antioxidant and cytoprotective protein genes, including enzymes that catalyze glutathione synthesis (Qiang Ma, 2013; Nguyen, Nioi, & Pickett, 2009). Knockout of *Nrf2* in mice results in increased sensitivity to oxidative stress-inducing agents including environmental toxicants, cigarette smoke, drugs and diseases(K. Chan, Han, & Kan, 2001; Cho, Reddy, & Kleeberger, 2006; Johnson, Amirahmadi, Ward, Fabry, & Johnson, 2009; Kensler, Wakabayashi, & Biswal, 2007; Q. Ma & He, 2012; Motohashi & Yamamoto, 2004; Rangasamy et al., 2004), while boosting

NRF2 activity has a protective effect against oxidative damage(Talalay, Dinkova-Kostova, & Holtzclaw, 2003). Nevertheless, mice lacking NRF2 have been shown to compensate for the deficiency by upregulating endothelial nitric oxide synthase (eNOS), which mediates cardioprotection against myocardia ischemia in conditions of decreased antioxidant capacity(Erkens et al., 2018). eNOS synthesizes nitric oxide, which has potent antioxidant effects(Erkens et al., 2018; Wink et al., 2001). Despite severely decreased glutathione levels and general antioxidant capacity, NRF2^-/-^ mice were able to preserve their redox status, vascular function, and resist myocardial ischemia/perfusion damage(Erkens et al., 2018). Nrf2 deficiency has also been shown to play a protective role against cardiomyopathy with molecular reprogramming of antioxidants at the transcript and protein levels playing a contributing role, with a noted increase in GPX1 transcript in NRF2 deficient mice with cardiomyopathy(Kannan et al., 2013). NRF2 and selenoproteins have been shown to work in conjunction to protect against oxidative damage(Kawatani, Suzuki, Shimizu, Kelly, & Yamamoto, 2011).

There are 25 Sec-containing proteins in humans, and 24 in rodents, with many serving as antioxidants in various thiol-dependent reactions (Hawkes & Alkan, 2010; Reeves & Hoffmann, 2009). Mice deficient in selenoprotein TRXR1, Selenoprotein R (SELR), and Selenoprotein P (SELP) have growth delays and display markers of oxidative stress and metabolic deficiencies (Bondareva et al., 2007; Fomenko et al., 2009; Schomburg et al., 2003). There are strong links between selenoprotein defects and inflammation. For example, selenoprotein S is an important regulator of the inflammatory response (Curran et al., 2005; Gao et al., 2006), with polymorphisms in the promoter region of *SelS* linked to decreased SELS expression and increased IL-6, IL-1β, and TNF-α levels (Curran et al., 2005). A deficiency or mutation in SELS is also associated with inflammation-related disorders and has been linked to gastric cancer (Mao, Cui, & Wang, 2015) and inflammatory bowel disease (Seiderer et al., 2007). In our study, we found a significant difference in SELS protein levels between WT and *Alkbh8^def^* NA treated mice, which may also explain the dramatic increase in IL-6 and other inflammatory markers at the transcript level in the *Alkbh8^def^* lungs. The dramatic increase in inflammation markers in writer deficient lungs can be also linked to increased ROS and decreased antioxidant enzymes in the form of specific seleno- and other proteins. In support of this idea, we observed deficiencies in selenoproteins TRXR1 and TRXR2 and non-selenoprotein TRX2 in *Alkbh8^de^*^f^ mice and associated increase in oxidative stress markers ORP and 8-isoprostanes. TRXs primarily reduce oxidized protein cysteine residues to form reduced disulfide bonds, with TRX2 localized to the mitochondria. TRXs are maintained in their reduced and active state by TRXRs using NADPH as a reducing equivalent, with TRXR1 located in the cytoplasm and TRXR2 in the mitochondria. This system is critical for signaling pathways involved in the protection from oxidants (Saccoccia et al., 2014). TRXRs are also required for development as complete knockout of TRXR1 and TRXR2 in mice result in severe growth abnormality and embryonic death at day 8.5 and 13, respectively(Bondareva et al., 2007; Conrad et al., 2004; Jakupoglu et al., 2005).

GPX family members catalyze the reduction of harmful peroxides by glutathione and play an essential role in protecting cells against oxidative damage. GPX1 is the most abundant selenoprotein in mice, and its importance in NA stress response is demonstrated by WT mice significantly upregulating GPX1 protein levels after NA exposure, whereas *Alkbh8^de^*^f^ mice fail to do so. *Alkbh8^de^*^f^ mice have decreased GPX1 and GPX3 levels compared to their carrier-treated counterparts. Previous studies have shown that glutathione (GSH) protects cells against NA metabolites (i.e. 1,2-naphthoquinone). GSH is a key detoxification pathway and its depletion is a major determinant of NA respiratory toxicity (Phimister et al., 2004; Plopper et al., 2013b). Many of the selenoproteins linked to ALKBH8 use GSH, including GPXs 1-6 in humans, and 1-5 in rodents, to transform H_2_O_2_ to water and lipid peroxides to alcohols. While the writer deficient lungs reprogram and upregulate other stress response systems under normal growth conditions, they are ill equipped to handle environmental challenge. These observations support the idea that dysregulated epitranscriptomic systems sensitizes lungs to NA-induced injury.

In general, cells and tissues respond to environmental stress by regulating stress response pathways. Lungs respond to toxicant challenge via recruitment of inflammatory cells and secretion of inflammatory cytokines, as well oxidative modification of biomolecules (in part due to high oxygen exposure in the lung and high surface contact with the environment). Oxidatively modified compounds, as well as bioactivated components of some environmental toxicants, are able to enter the blood stream and other organs and produce systemic oxidative stress and inflammation (Dalle-Donne, Rossi, Colombo, Giustarini, & Milzani, 2006; Gomez-Mejiba et al., 2009). Thus, inhaled environmental toxicants often have an effect that exceeds that of the lung. We have observed this phenomime in our *Alkbh8^def^* mice, as weigh loss is increased and animal survival decreased after NA-exposure, which is likely due to systemic issues. Our study demonstrates that the epitranscriptomic writer ALKBH8 is required to mitigate the effects of NA. It represents one of the first examples of how a mammalian RNA modification system is vital after environmental challenge. As there are over twenty epitranscriptomic writers in mammals(Kadumuri & Janga, 2018), there are most likely other RNA modifications, writers and perhaps erasers essential to the response to common environmental toxicants.

The highest occupational exposure to naphthalene occurs in industries dealing with wood treatment, coal tar production, mothball production and jet fuel industries. NA bioactivation by CYP enzymes is a well-documented requirement of NA toxicity (ATSDR, 2005; L. Li et al., 2011). Bioactivation and now regulators of selenoproteins, in this case ALKBH8-dependent epitranscriptomic control, should also be considered modifiers of NA exposure effects. Age-associated decline in GSH synthesis and activity has been reported in mice and rats (Chen, Richie, & Lang, 1989; Kim et al., 2003; Liu, 2002). Deceased ALKBH8 has been identified in aged fibroblasts, relative to young, with deficiencies also linked to early senescence(Lee et al., 2019). Due to age induced changes in transcription and epitranscriptomic activity, NA sensitivity may be amplified in older people, but that is speculation that needs to be further studied. In support though, age related decreases have been established for drug and xenobiotic metabolism capacity in aging mice(Kwak, Kim, Oh, & Kim, 2015) and humans (Sotaniemi, Arranto, Pelkonen, & Pasanen, 1997).

Tolerance is defined as resistance to a high challenging dose after repeated exposures. Upregulation of glutathione synthesis has been reported as a tolerance mechanism in lung airway epithelial club cells, (Plopper et al., 2013b; Van Winkle et al., 1995; West et al., 2003). Glutathione depletion is a hallmark of NA toxicity (Phimister et al., 2004). γ-GCS is the rate-limiting enzyme in GSH synthesis, and treatment of mice with buthionine sulfoximine, a γ-GCS inhibitor, eliminates the tolerant phenotype in a majority of mice post-NA exposure (West et al., 2003, 2002). Selenoprotein H (SELH) has been shown to be an important transcription factor promoting expression of genes involved in GSH synthesis, including γ-GCS (Panee et al., 2007). Decreased SELH may explain our finding that ALKBH8-deficient mice fail to develop tolerance to NA, but we were unable to identify suitable antibodies to measure it. Our study highlights the importance of ALKBH8 in promoting tolerance, most likely via translational regulation of selenoproteins and GSH levels. Dysregulated transcriptional response due to the molecular reprogramming that occurs in response to the epitranscriptomic deficiency may also prevent tolerance in the *Alkbh8^def^* mice. It is notable though that tolerance to a high dose chemical challenge has been demonstrated for a homolog of ALKBH8, namely *E. coli* AlkB (Samson & Cairns, 1977). The adaptive response to alkylating agents was characterized under conditions that mimic tolerance, as *E.coli* cells pretreated with a low dose of alkylating agent were able to survive a high dose challenge, relative to naive cells. The adaptive response includes up-regulation of members of the Ada operon, the DNA glycosylase AlkA and the demethylase AlkB. A major finding of our study is the determination that the epitranscriptomic writer ALKBH8 is a key driver of tolerance, which highlights a novel role for tRNA modifications and translational regulatory mechanisms in environmental responses.

## Conclusions

Our study supports a model (Fig. 7) in which ALKBH8-catalyzed epitranscriptomic marks promote the translation of selenoproteins important for handling stress, preventing NA toxicity and driving tolerance. Efforts to identify other epitranscriptomic writers and marks, and translational control systems that mammals use to survive and tolerate NA and other exposures, are needed to further understand the roles of RNA modifications in environmental exposure induced pathobiological changes.

**Figure 7.**
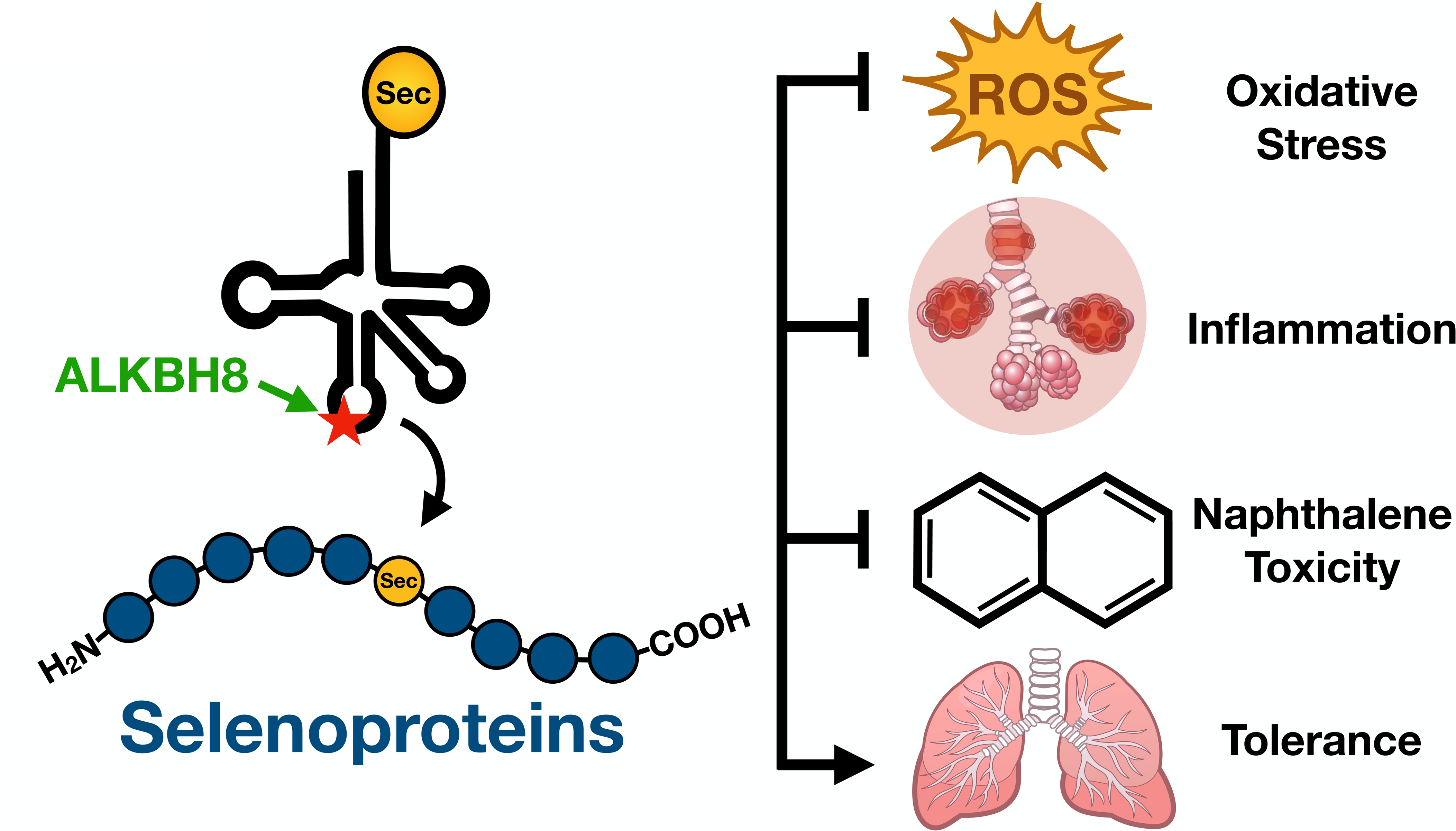
Model. Our study supports a model in which ALKBH8-catalyzed epitranscriptomic marks promote the translation of selenoproteins important for handling stress, preventing NA toxicity and driving tolerance.

## Acknowledgements

Research was funded in part by grants from the National Institutes of Health (R01ES020867, P30ES006694, R01ES026856 and R01ES024615). We would like to thank our colleagues who have provided helpful suggestions. We would also like to thank the dedicated animal care staff at the University and Albany, Wadsworth Institute, and University of Arizona for their helpful assistance in all issues related to our animal colonies and Ms. Weizhu Yang for assistance with mouse breeding.

