## Supplemental Materials for "Epitranscriptomic regulation of the response to the air pollutant naphthalene in mouse lungs: from the perspectives of specialized translation and tolerance linked to the writer ALKBH8"

### Supplemental Figure 1.

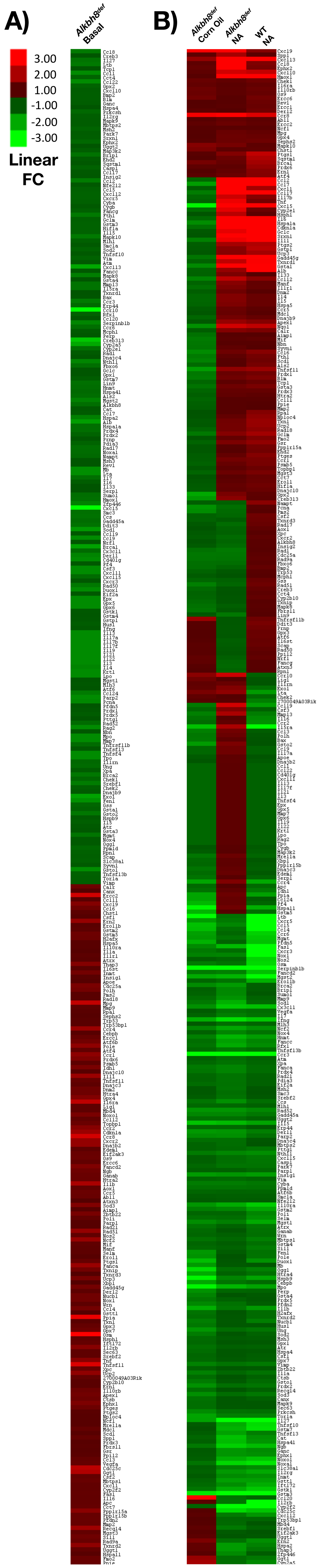

Supplemental Figure 2.

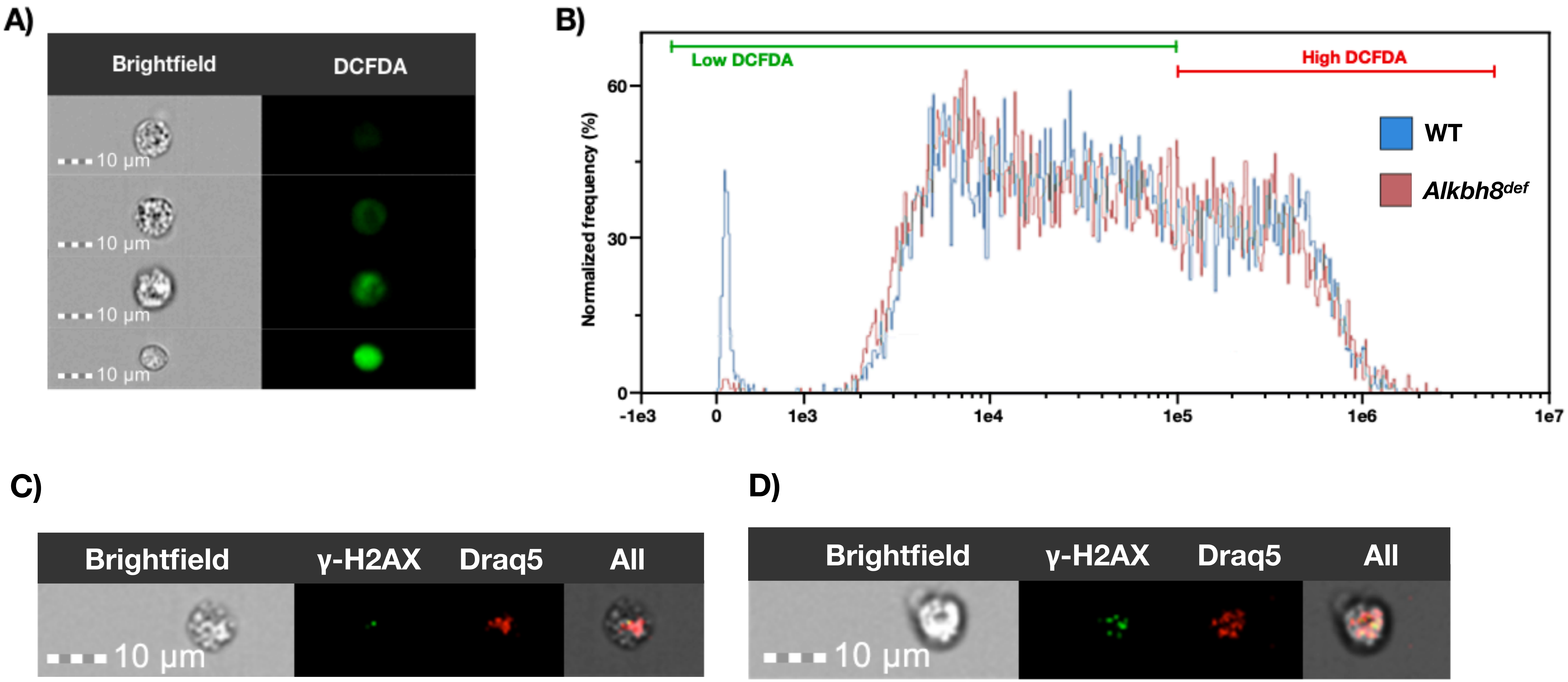

#### Supplemental Figure 3.

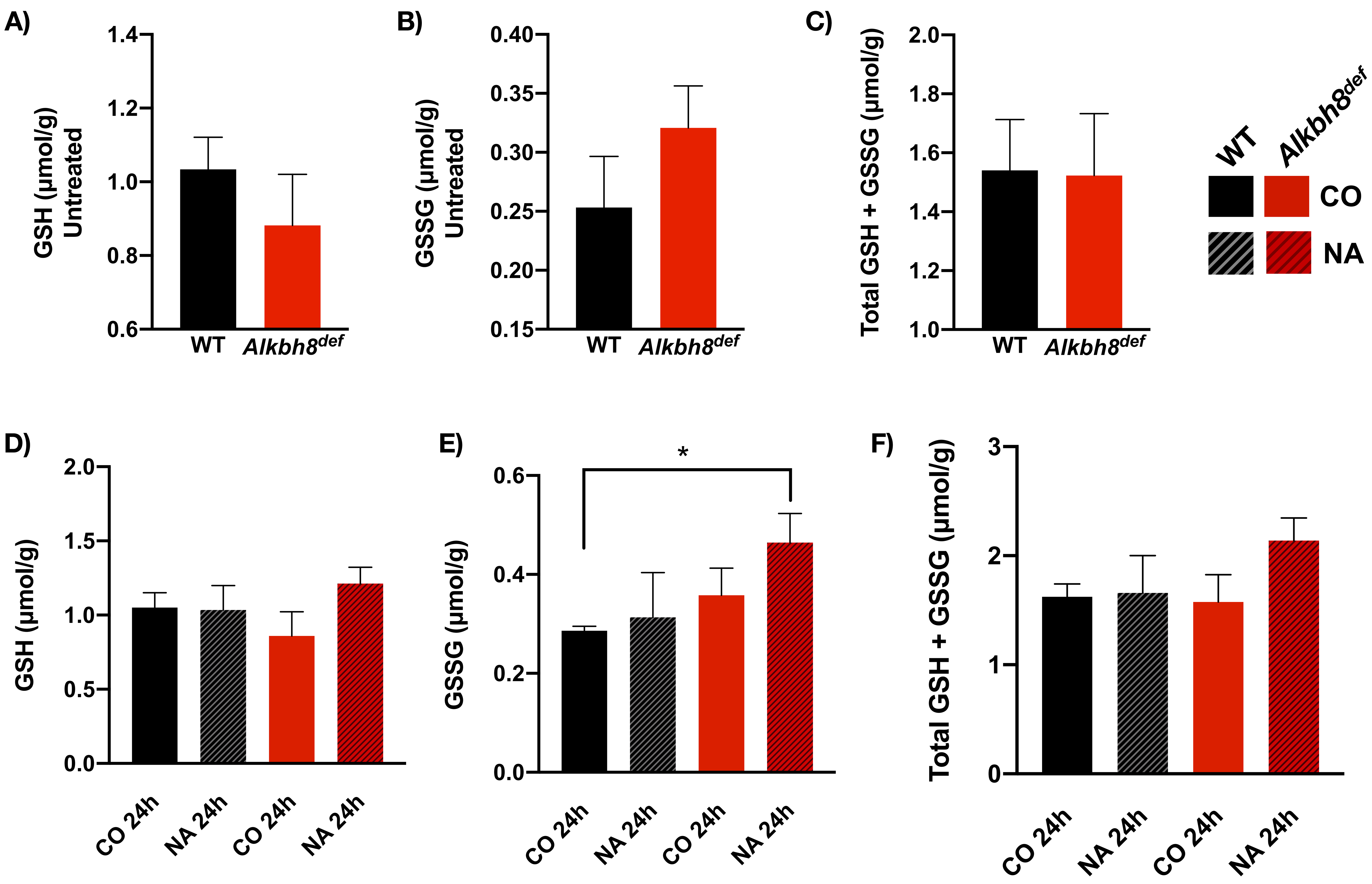

**Supplemental Figure 4.**

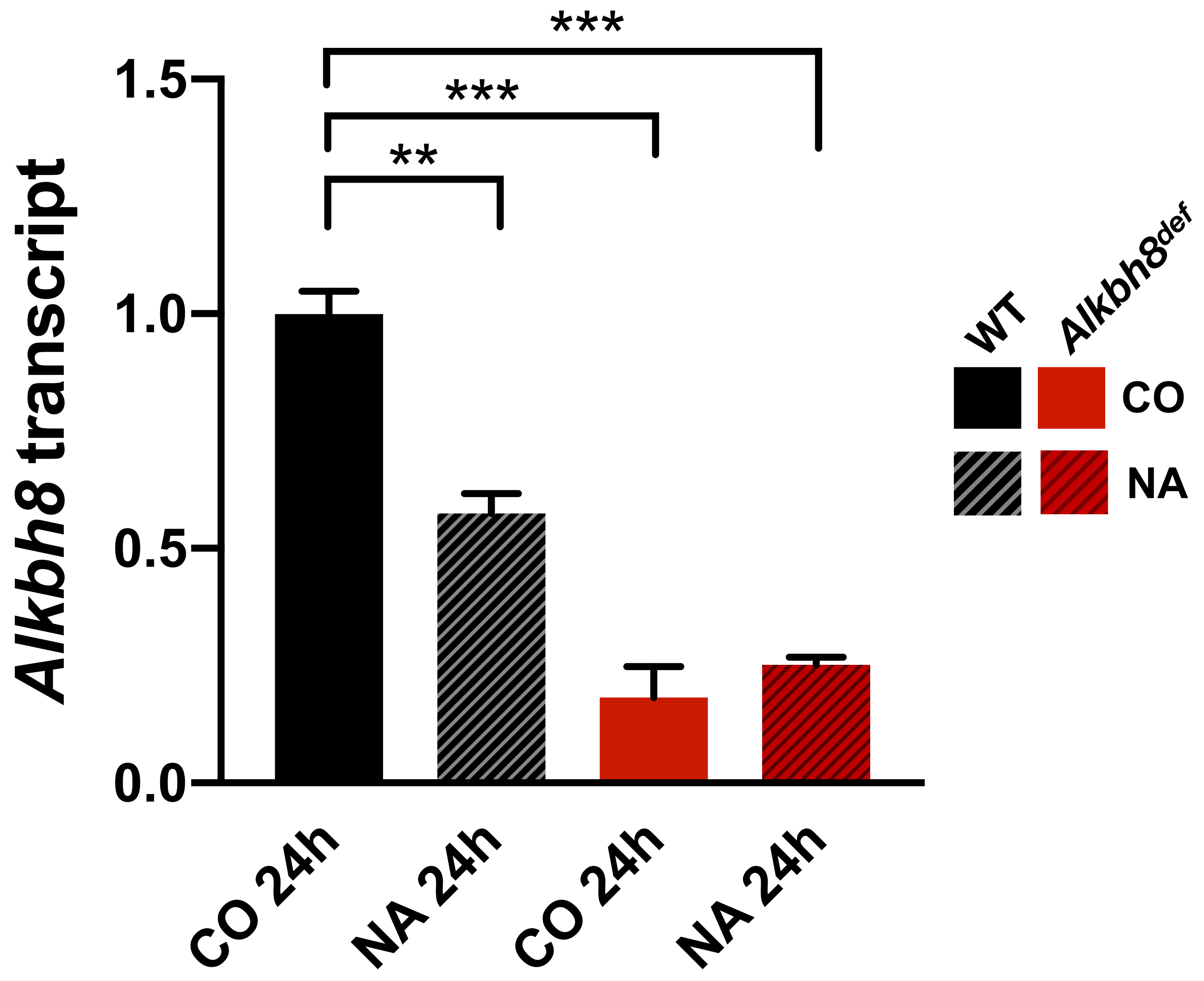

**Supplemental Figure 5.**

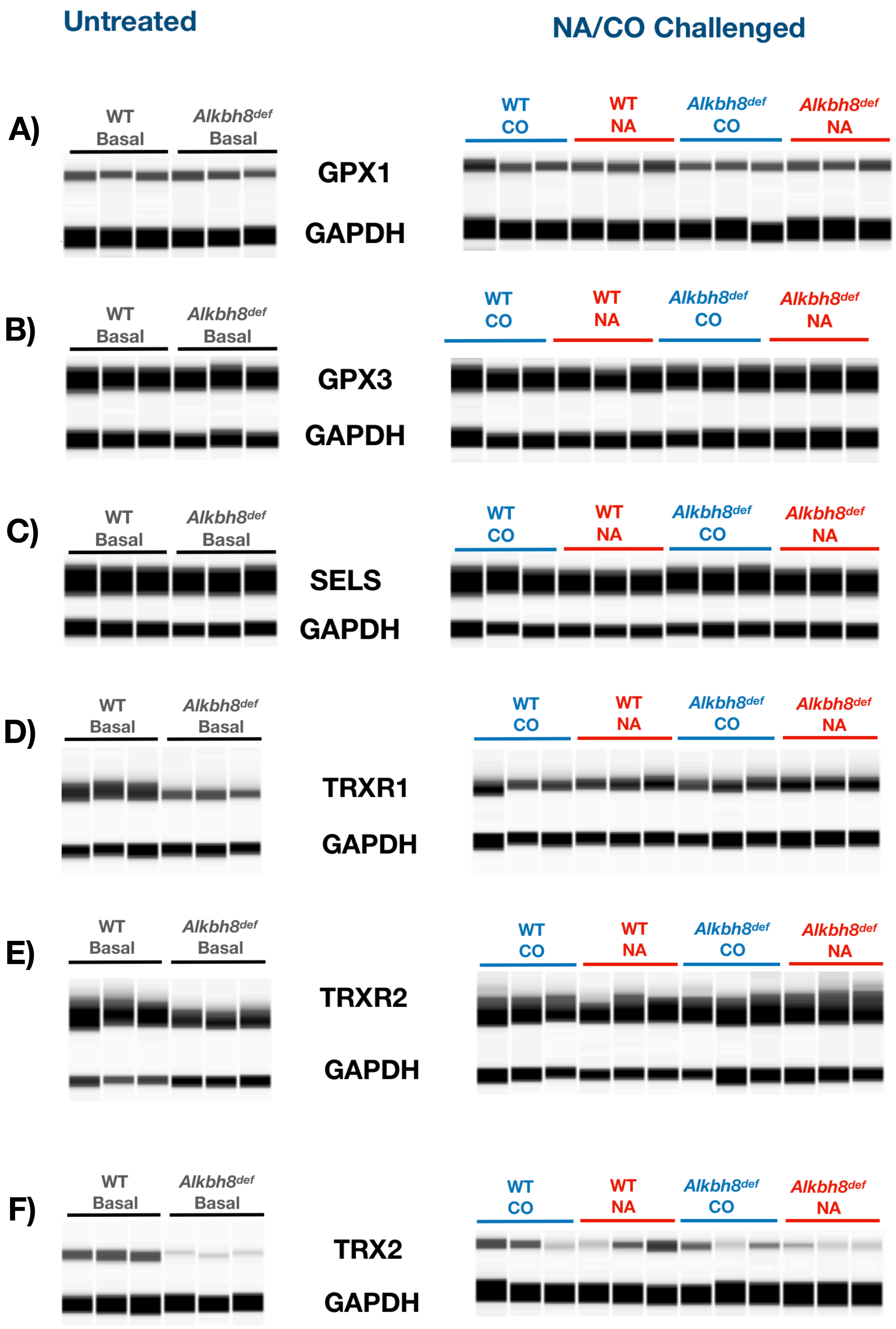

**Supplemental Figure 6.**

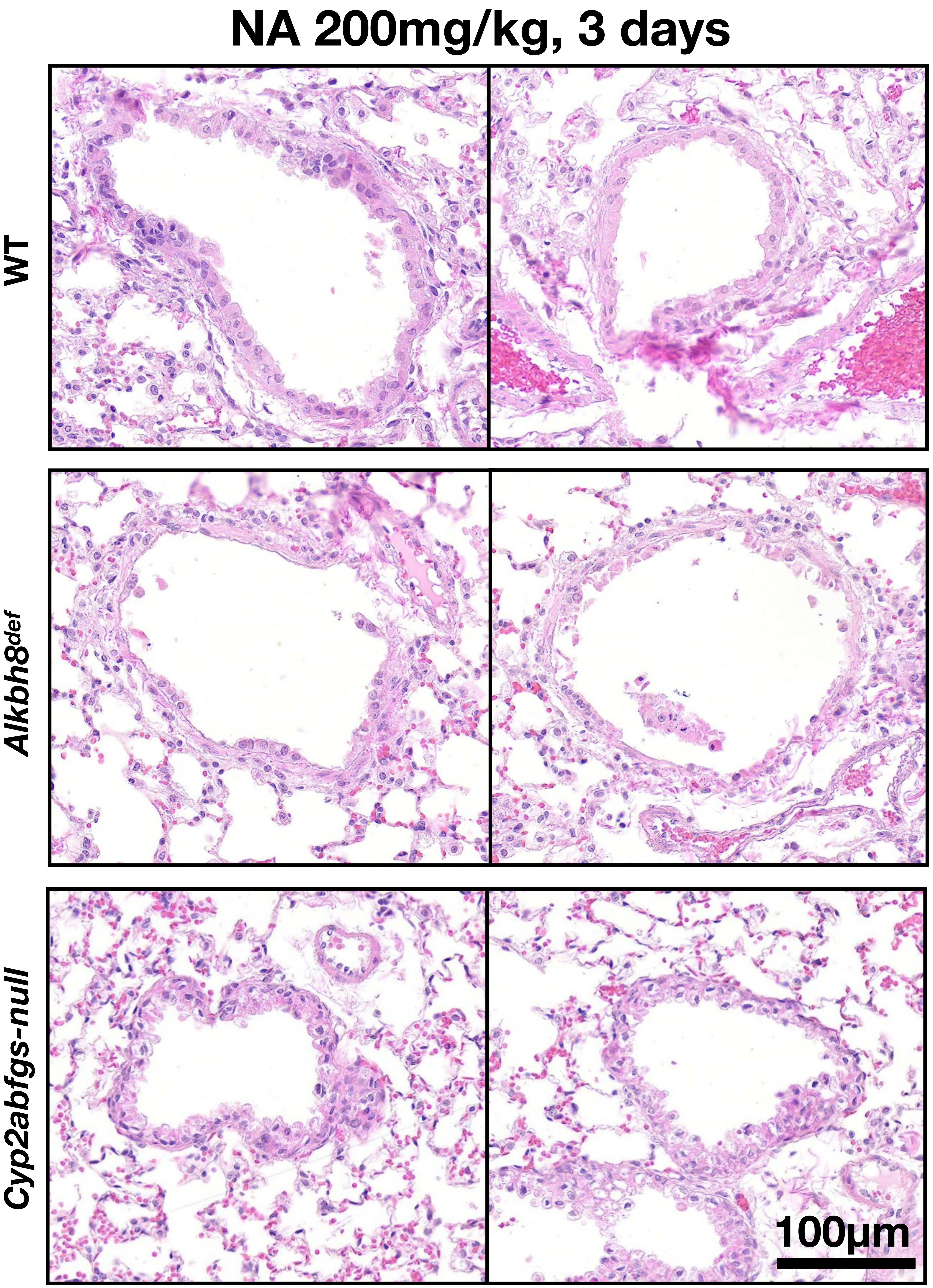

**Supplemental Table 1: Targeted RNAseq Gene Panel**

| Genes on panel: | Ensembl ID: |
|---|---|
| *Abl1* | ENSMUSG00000026842 |
| *Aimp1* | ENSMUSG00000028029 |
| *Alb* | ENSMUSG00000029368 |
| *Als2* | ENSMUSG00000026024 |
| *Aox1* | ENSMUSG00000063558 |
| *Apc* | ENSMUSG00000005871 |
| *Apex1* | ENSMUSG00000035960 |
| *Apoe* | ENSMUSG00000002985 |
| *Atf4* | ENSMUSG00000042406 |
| *Atf6* | ENSMUSG00000026663 |
| *Atf6b* | ENSMUSG00000015461 |
| *Atm* | ENSMUSG00000034218 |
| *Atr* | ENSMUSG00000032409 |
| *Atrx* | ENSMUSG00000031229 |
| *Atxn3* | ENSMUSG00000021189 |
| *Bax* | ENSMUSG00000003873 |
| *Blm* | ENSMUSG00000030528 |
| *Bmp2* | ENSMUSG00000027358 |
| *Brca1* | ENSMUSG00000017146 |
| *Brca2* | ENSMUSG00000041147 |
| *Brip1* | ENSMUSG00000034329 |
| *Calr* | ENSMUSG00000003814 |
| *Canx* | ENSMUSG00000020368 |
| *Casp1* | ENSMUSG00000025888 |
| *Cat* | ENSMUSG00000027187 |
| *Ccl1* | ENSMUSG00000020702 |
| *Ccl11* | ENSMUSG00000020676 |
| *Ccl12* | ENSMUSG00000035352 |
| *Ccl17* | ENSMUSG00000031780 |
| *Ccl19* | ENSMUSG00000071005 |
| *Ccl2* | ENSMUSG00000035385 |
| *Ccl20* | ENSMUSG00000026166 |
| *Ccl22* | ENSMUSG00000031779 |
| *Ccl24* | ENSMUSG00000004814 |
| *Ccl3* | ENSMUSG00000000982 |
| *Ccl4* | ENSMUSG00000018930 |
| *Ccl5* | ENSMUSG00000035042 |

| | |
|---|---|
| *Ccl6* | ENSMUSG00000018927 |
| *Ccl7* | ENSMUSG00000035373 |
| *Ccl8* | ENSMUSG00000009185 |
| *Ccl9* | ENSMUSG00000019122 |
| *Ccr1* | ENSMUSG00000025804 |
| *Ccr10* | ENSMUSG00000044052 |
| *Ccr2* | ENSMUSG00000049103 |
| *Ccr3* | ENSMUSG00000035448 |
| *Ccr4* | ENSMUSG00000047898 |
| *Ccr5* | ENSMUSG00000079227 |
| *Ccr6* | ENSMUSG00000040899 |
| *Ccr8* | ENSMUSG00000042262 |
| *Ccs* | ENSMUSG00000034108 |
| *Cct4* | ENSMUSG00000007739 |
| *Cct7* | ENSMUSG00000030007 |
| *Cd40lg* | ENSMUSG00000031132 |
| *Cdc25a* | ENSMUSG00000032477 |
| *Cdc25c* | ENSMUSG00000044201 |
| *Cdkn1a* | ENSMUSG00000023067 |
| *Cebpb* | ENSMUSG00000056501 |
| *Chek1* | ENSMUSG00000032113 |
| *Chek2* | ENSMUSG00000029521 |
| *Chst1* | ENSMUSG00000027221 |
| *Creb3* | ENSMUSG00000028466 |
| *Creb3l3* | ENSMUSG00000035041 |
| *Csf1* | ENSMUSG00000014599 |
| *Csf2* | ENSMUSG00000018916 |
| *Csf3* | ENSMUSG00000038067 |
| *Ctsb* | ENSMUSG00000021939 |
| *Cx3cl1* | ENSMUSG00000031778 |
| *Cxcl1* | ENSMUSG00000029380 |
| *Cxcl10* | ENSMUSG00000034855 |
| *Cxcl11* | ENSMUSG00000060183 |
| *Cxcl12* | ENSMUSG00000061353 |
| *Cxcl13* | ENSMUSG00000023078 |
| *Cxcl15* | ENSMUSG00000029375 |
| *Cxcl5* | ENSMUSG00000029371 |
| *Cxcl9* | ENSMUSG00000029417 |
| *Cxcr2* | ENSMUSG00000026180 |
| *Cxcr3* | ENSMUSG00000050232 |
| *Cxcr5* | ENSMUSG00000047880 |

| | |
|---|---|
| *Cyba* | ENSMUSG00000006519 |
| *Cygb* | ENSMUSG00000020810 |
| *Cyp2a5* | ENSMUSG00000005547 |
| *Cyp2b10* | ENSMUSG00000030483 |
| *Cyp2e1* | ENSMUSG00000025479 |
| *Cyp2f2* | ENSMUSG00000052974 |
| *Ddit3* | ENSMUSG00000025408 |
| *Derl1* | ENSMUSG00000022365 |
| *Derl2* | ENSMUSG00000018442 |
| *Dnajb2* | ENSMUSG00000026203 |
| *Dnajb9* | ENSMUSG00000014905 |
| *Dnajc10* | ENSMUSG00000027006 |
| *Dnajc3b* | ENSMUSG00000022136 |
| *Dnajc4* | ENSMUSG00000024963 |
| *Dnm2* | ENSMUSG00000033335 |
| *Duox1* | ENSMUSG00000033268 |
| *Edem1* | ENSMUSG00000030104 |
| *Ehd2* | ENSMUSG00000074364 |
| *Eif2a* | ENSMUSG00000027810 |
| *Eif2ak3* | ENSMUSG00000031668 |
| *Ephx1* | ENSMUSG00000038776 |
| *Ephx2* | ENSMUSG00000022040 |
| *Epx* | ENSMUSG00000052234 |
| *Ercc1* | ENSMUSG00000003549 |
| *Ercc2* | ENSMUSG00000030400 |
| *Ercc6* | ENSMUSG00000054051 |
| *Ern1* | ENSMUSG00000020715 |
| *Ern2* | ENSMUSG00000030866 |
| *Ero1l* | ENSMUSG00000021831 |
| *Ero1lb* | ENSMUSG00000057069 |
| *Erp44* | ENSMUSG00000028343 |
| *Exo1* | ENSMUSG00000039748 |
| *Fanca* | ENSMUSG00000032815 |
| *Fancc* | ENSMUSG00000021461 |
| *Fancd2* | ENSMUSG00000034023 |
| *Fancg* | ENSMUSG00000028453 |
| *Fasl* | ENSMUSG00000000817 |
| *Fbxo6* | ENSMUSG00000055401 |
| *Fen1* | ENSMUSG00000024742 |
| *Fmo2* | ENSMUSG00000040170 |
| *Fth1* | ENSMUSG00000024661 |

| | |
|---|---|
| *Gadd45a* | ENSMUSG00000036390 |
| *Gadd45g* | ENSMUSG00000021453 |
| *Ganab* | ENSMUSG00000071650 |
| *Ganc* | ENSMUSG00000062646 |
| *Gclc* | ENSMUSG00000032350 |
| *Gclm* | ENSMUSG00000028124 |
| *Ggt1* | ENSMUSG00000006345 |
| *Gpx1* | ENSMUSG00000063856 |
| *Gpx2* | ENSMUSG00000042808 |
| *Gpx3* | ENSMUSG00000018339 |
| *Gpx4* | ENSMUSG00000075706 |
| *Gpx5* | ENSMUSG00000004344 |
| *Gpx6* | ENSMUSG00000004341 |
| *Gpx7* | ENSMUSG00000028597 |
| *Gsr* | ENSMUSG00000031584 |
| *Gss* | ENSMUSG00000027610 |
| *Gsta1* | ENSMUSG00000074183 |
| *Gsta3* | ENSMUSG00000025934 |
| *Gsta4* | ENSMUSG00000032348 |
| *Alkbh8* | ENSMUSG00000025899 |
| *Gstk1* | ENSMUSG00000029864 |
| *Gstm2* | ENSMUSG00000040562 |
| *Gstm3* | ENSMUSG00000004038 |
| *Gstm4* | ENSMUSG00000027890 |
| *Gstm5* | ENSMUSG00000004032 |
| *Gstm7* | ENSMUSG00000004035 |
| *Gsto1* | ENSMUSG00000025068 |
| *Gsto2* | ENSMUSG00000025069 |
| *Gstp1* | ENSMUSG00000060803 |
| *Gstt1* | ENSMUSG00000001663 |
| *H2afx* | ENSMUSG00000049932 |
| *Hif1a* | ENSMUSG00000021109 |
| *Hmox1* | ENSMUSG00000005413 |
| *Hnmt* | ENSMUSG00000026986 |
| *Hspa1a* | ENSMUSG00000091971 |
| *Hspa1l* | ENSMUSG00000007033 |
| *Hspa2* | ENSMUSG00000059970 |
| *Hspa4* | ENSMUSG00000020361 |
| *Hspa4l* | ENSMUSG00000025757 |
| *Hspa5* | ENSMUSG00000026864 |
| *Hspb9* | ENSMUSG00000017832 |

| | |
|---|---|
| *Hsph1* | ENSMUSG00000029657 |
| *Htra2* | ENSMUSG00000068329 |
| *Htra4* | ENSMUSG00000037406 |
| *Hus1* | ENSMUSG00000020413 |
| *Idh1* | ENSMUSG00000025950 |
| *Ifng* | ENSMUSG00000055170 |
| *Ift172* | ENSMUSG00000038564 |
| *Il10ra* | ENSMUSG00000032089 |
| *Il10rb* | ENSMUSG00000022969 |
| *Il11* | ENSMUSG00000004371 |
| *Il13* | ENSMUSG00000020383 |
| *Il15* | ENSMUSG00000031712 |
| *Il16* | ENSMUSG00000001741 |
| *Il17a* | ENSMUSG00000025929 |
| *Il17b* | ENSMUSG00000024578 |
| *Il17f* | ENSMUSG00000041872 |
| *Il19* | ENSMUSG00000016524 |
| *Il1a* | ENSMUSG00000027399 |
| *Il1b* | ENSMUSG00000027398 |
| *Il1r1* | ENSMUSG00000026072 |
| *Il1rn* | ENSMUSG00000026981 |
| *Il21* | ENSMUSG00000027718 |
| *Il22* | ENSMUSG00000074695 |
| *Il27* | ENSMUSG00000044701 |
| *Il2rb* | ENSMUSG00000068227 |
| *Il2rg* | ENSMUSG00000031304 |
| *Il3* | ENSMUSG00000018914 |
| *Il33* | ENSMUSG00000024810 |
| *Il4* | ENSMUSG00000000869 |
| *Il5* | ENSMUSG00000036117 |
| *Il5ra* | ENSMUSG00000005364 |
| *Il6* | ENSMUSG00000025746 |
| *Il6ra* | ENSMUSG00000027947 |
| *Il6st* | ENSMUSG00000021756 |
| *Il7* | ENSMUSG00000040329 |
| *Inmt* | ENSMUSG00000003477 |
| *Insig1* | ENSMUSG00000045294 |
| *Insig2* | ENSMUSG00000003721 |
| *Krt1* | ENSMUSG00000046834 |
| *Lig1* | ENSMUSG00000056394 |
| *Lin9* | ENSMUSG00000058729 |

| | |
|---|---|
| *Lpo* | ENSMUSG00000009356 |
| *Lta* | ENSMUSG00000024402 |
| *Ltb* | ENSMUSG00000024399 |
| *Manf* | ENSMUSG00000032575 |
| *Mapk10* | ENSMUSG00000046709 |
| *Mapk8* | ENSMUSG00000021936 |
| *AW490526* | ENSMUSG00000020366 |
| *Mb* | ENSMUSG00000018893 |
| *Mbd4* | ENSMUSG00000030322 |
| *Mbtps1* | ENSMUSG00000031835 |
| *Mbtps2* | ENSMUSG00000046873 |
| *Mcph1* | ENSMUSG00000039842 |
| *Mdc1* | ENSMUSG00000061607 |
| *Mgmt* | ENSMUSG00000054612 |
| *Mgst1* | ENSMUSG00000008540 |
| *Mgst2* | ENSMUSG00000074604 |
| *Mgst3* | ENSMUSG00000026688 |
| *Mif* | ENSMUSG00000033307 |
| *Mlh1* | ENSMUSG00000032498 |
| *Mlh3* | ENSMUSG00000021245 |
| *Mmp13* | ENSMUSG00000050578 |
| *Mmp2* | ENSMUSG00000031740 |
| *Mmp7* | ENSMUSG00000018623 |
| *Mmp9* | ENSMUSG00000017737 |
| *Mpg* | ENSMUSG00000020287 |
| *Mpo* | ENSMUSG00000009350 |
| *Mre11a* | ENSMUSG00000031928 |
| *Msh2* | ENSMUSG00000024151 |
| *Msh3* | ENSMUSG00000014850 |
| *Nampt* | ENSMUSG00000020572 |
| *Nbn* | ENSMUSG00000028224 |
| *Ncf1* | ENSMUSG00000015950 |
| *Ncf2* | ENSMUSG00000026480 |
| *Nfe2l2* | ENSMUSG00000015839 |
| *Nrf1* | ENSMUSG00000058440 |
| *Ngb* | ENSMUSG00000021032 |
| *Nos2* | ENSMUSG00000020826 |
| *Nox1* | ENSMUSG00000031257 |
| *Nox4* | ENSMUSG00000030562 |
| *Noxa1* | ENSMUSG00000036805 |
| *Noxo1* | ENSMUSG00000019320 |

| | |
|---|---|
| *Nploc4* | ENSMUSG00000039703 |
| *Nqo1* | ENSMUSG00000003849 |
| *Nthl1* | ENSMUSG00000041429 |
| *Nucb1* | ENSMUSG00000030824 |
| *Ogg1* | ENSMUSG00000030271 |
| *Os9* | ENSMUSG00000040462 |
| *Osm* | ENSMUSG00000058755 |
| *Park7* | ENSMUSG00000028964 |
| *Parp1* | ENSMUSG00000026496 |
| *Parp2* | ENSMUSG00000036023 |
| *Pcna* | ENSMUSG00000027342 |
| *Pdia3* | ENSMUSG00000027248 |
| *Perp* | ENSMUSG00000019851 |
| *Pf4* | ENSMUSG00000029373 |
| *Pfdn2* | ENSMUSG00000006412 |
| *Pfdn5* | ENSMUSG00000001289 |
| *Pms2* | ENSMUSG00000079109 |
| *Pole* | ENSMUSG00000007080 |
| *Polh* | ENSMUSG00000023953 |
| *Poli* | ENSMUSG00000038425 |
| *POR* | ENSMUSG00000005514 |
| *Ppia* | ENSMUSG00000071866 |
| *Ppm1d* | ENSMUSG00000020525 |
| *Ppp1r15a* | ENSMUSG00000040435 |
| *Ppp1r15b* | ENSMUSG00000046062 |
| *Prdx1* | ENSMUSG00000028691 |
| *Prdx2* | ENSMUSG00000005161 |
| *Prdx3* | ENSMUSG00000024997 |
| *Prdx4* | ENSMUSG00000025289 |
| *Prdx5* | ENSMUSG00000024953 |
| *Prdx6* | ENSMUSG00000026701 |
| *Prkcsh* | ENSMUSG00000003402 |
| *Prnp* | ENSMUSG00000079037 |
| *Psmb5* | ENSMUSG00000022193 |
| *Ptges* | ENSMUSG00000050737 |
| *Ptgs1* | ENSMUSG00000047250 |
| *Ptgs2* | ENSMUSG00000032487 |
| *Pttg1* | ENSMUSG00000020415 |
| *Rad1* | ENSMUSG00000022248 |
| *Rad17* | ENSMUSG00000021635 |
| *Rad18* | ENSMUSG00000030254 |

| | |
|---|---|
| *Rad21* | ENSMUSG00000022314 |
| *Rad50* | ENSMUSG00000020380 |
| *Rad51* | ENSMUSG00000027323 |
| *Rad52* | ENSMUSG00000030166 |
| *Rad9a* | ENSMUSG00000024824 |
| *Rag2* | ENSMUSG00000032864 |
| *Recql4* | ENSMUSG00000033762 |
| *Rev1* | ENSMUSG00000026082 |
| *Rpa1* | ENSMUSG00000000751 |
| *Rpn1* | ENSMUSG00000030062 |
| *Scap* | ENSMUSG00000032485 |
| *Scd1* | ENSMUSG00000037071 |
| *Sec63* | ENSMUSG00000019802 |
| *Selm* | ENSMUSG00000075702 |
| *Selp* | ENSMUSG00000026580 |
| *SepN1* | ENSMUSG00000050989 |
| *Sephs2* | ENSMUSG00000049091 |
| *Serp1* | ENSMUSG00000027808 |
| *Serpinb1b* | ENSMUSG00000051029 |
| *Sil1* | ENSMUSG00000024357 |
| *Slc38a1* | ENSMUSG00000023169 |
| *Smc1a* | ENSMUSG00000041133 |
| *Smc3* | ENSMUSG00000024974 |
| *Sod1* | ENSMUSG00000022982 |
| *Sod2* | ENSMUSG00000006818 |
| *Sod3* | ENSMUSG00000072941 |
| *Spp1* | ENSMUSG00000029304 |
| *Sqstm1* | ENSMUSG00000015837 |
| *Srebf1* | ENSMUSG00000020538 |
| *Srebf2* | ENSMUSG00000022463 |
| *Srxn1* | ENSMUSG00000032802 |
| *Sumo1* | ENSMUSG00000026021 |
| *Syvn1* | ENSMUSG00000024807 |
| *Tcp1* | ENSMUSG00000068039 |
| *Tnf* | ENSMUSG00000024401 |
| *AI315324* | ENSMUSG00000063727 |
| *Tnfsf10* | ENSMUSG00000039304 |
| *Tnfsf11* | ENSMUSG00000022015 |
| *Tnfsf13* | ENSMUSG00000089669 |
| *Tnfsf13b* | ENSMUSG00000031497 |
| *Tnfsf4* | ENSMUSG00000026700 |

| | |
|---|---|
| *Topbp1* | ENSMUSG00000032555 |
| *Tor1a* | ENSMUSG00000026849 |
| *Tpo* | ENSMUSG00000020673 |
| *Trp53* | ENSMUSG00000059552 |
| *Trp53bp1* | ENSMUSG00000043909 |
| *Txn1* | ENSMUSG00000028367 |
| *Txnip* | ENSMUSG00000038393 |
| *Txnrd1* | ENSMUSG00000020250 |
| *Txnrd2* | ENSMUSG00000075704 |
| *Txnrd3* | ENSMUSG00000000811 |
| *Ucp2* | ENSMUSG00000033685 |
| *Ucp3* | ENSMUSG00000032942 |
| *Uggt1* | ENSMUSG00000037470 |
| *Uggt2* | ENSMUSG00000042104 |
| *Ung* | ENSMUSG00000029591 |
| *Vegfa* | ENSMUSG00000023951 |
| *Vim* | ENSMUSG00000026728 |
| *Vimp* | ENSMUSG00000075701 |
| *Wrn* | ENSMUSG00000031583 |
| *Xbp1* | ENSMUSG00000020484 |
| *Xpa* | ENSMUSG00000028329 |
| *Xpc* | ENSMUSG00000030094 |

**Supplemental Tables and Figure Legends**

**Supplemental Table 1. Complete gene panel for targeted RNA sequencing**

Gene names are shown on the left and Ensembl ID on the right.

**Supplemental Figure 1. Expression of 350 stress response transcripts obtained via targeted RNA sequencing.**

Heat map showing differences in quantitated gene expression of 350 stress response genes. **A)** Basal *Alkbh8*$^{def}$ lung transcripts relative to WT **B)** Corn Oil and 24 hour, 200mg/kg NA-treated WT and *Alkbh8*$^{def}$ lungs. N = 2 mice per condition.

**Supplemental Figure 2. Amnis Imagestream experimental data showing DCFDA results and γH2AX examples.**

Intracelluar ROS was measured in mouse lungs by DCFDA staining. Untreated 8-12 week old WT and *Alkbh8*$^{def}$ mice were sacrificed, lungs removed and enzymatically and mechanically dissociated into a single-cell suspension as described in methods. The live cells were then incubated with DCFDA dye and imaged with an Amnis Imagestream ISX100 flow cytometer. **A)** Representative images showing range of DCFDA fluorescence in the mixed population of lung cells. **B)** The frequency of WT (blue) and *Alkbh8*$^{def}$ (red) cells exhibiting specific DCFDA intensities is plotted for a population of 12,160 and 11,726, respectively. An arbitrary cutoff of $1x10^5$ RFU was used to separate low and high DCFDA intensity. 30.6% of WT cells fall into the high DCFDA category, compared to 30.9% of *Alkbh8^def^* (P = 0.946). γH2AX staining was performed using the Amnis Imagestream ISX100 flow cytometer. **C)** shows representative images of a low spot count ≤1 and **D)** high spot count ≥2.

**Supplemental Figure 3. *Alkbh8$^{def}$* mice have less GSH and more GSSG under basal conditions, and more GSSG under NA-challenged conditions.**

LC-MS/MS was employed to measure GSH and GSSG in WT and *Alkbh8$^{def}$* mice. **A)** GSH in untreated lungs **B)** GSSG in untreated lungs **C)** Total GSH + GSSG in untreated lungs **D)** GSH in CO and NA-challenged lungs and **E)** GSSG in CO and NA-challenged lungs **F)** Total GSH + GSSG in NA-challenged lungs. (N = 3 mice per condition, *P ≤ 0.05).

**Supplemental Figure 4. *Alkbh8$^{def}$* mice have less *Alkbh8* transcript after CO and NA challenge.**

Quantitative RT-PCR (qRT-PCR) was used to measure *Alkbh8* transcript in 24h CO and 24h, 200 mg/kg NA-treated mouse lungs as described in the methods. (N = 3 lungs per condition, **P ≤ 0.01, ***P ≤ 0.001).

**Supplemental Figure 5. Wes blots for selenoproteins GPX1, GPX3, SELS, TRXR1 and TRXR2 and non-selenoprotein TRX2.**

Wes capillaries showing full protein expression results for **A)** GPX1 **B)** GPX3, **C)** SELS **D)** TRXR1, **E**) TRXR2, and **F)** TRX2. Loading control (GAPDH) was multiplexed in each capillary and is shown on the bottom in each data set. Each capillary blot represents one mouse.

**Supplemental Figure 6. *Alkbh8$^{def}$* mice show increased lung injury after 3-day NA exposure.**

Two examples from lungs of WT (top row), *Alkbh8$^{def}$* (middle row) and *Cyp2abfgs-null* (bottom row) mouse lungs after 3-day NA exposure, 200 mg/kg.
